# Computational Model of Flower Pattern Evolution Predicts Spontaneous Emergence of Boundary Cell Types Across Petal Epidermis

**DOI:** 10.1101/2025.05.03.651954

**Authors:** Steven Oud, Maciej M. Żurowski, Pjotr L. van der Jagt, May T. S. Yeo, Joseph F. Walker, Edwige Moyroud, Renske M. A. Vroomans

**Affiliations:** The Sainsbury Laboratory, University of Cambridge, Bateman Street, Cambridge CB2 1LR, UK; Department of Genetics, Downing Site, University of Cambridge, Cambridge CB2 3EJ, UK; Department of Biological Sciences, University of Illinois at Chicago, Chicago, IL 60607, USA

**Keywords:** Evo-devo, Petal patterning, Gene regulatory networks, Gene expression noise, Boundary establishment, Cell fate

## Abstract

Petal patterns play an important role in the reproductive success of flowering plants by attracting pollinators and protecting reproductive organs from environmental factors. Some transcription factors (TFs) that control pigment production and cuticle elaboration in petal epidermal cells have been identified. However, little is known about the upstream developmental processes that pre-pattern the petal surface to first establish the different domains where these regulators will later be expressed. Here, we developed a computational model of the evolution and development of petal patterns to investigate this early pre-patterning phase. We selected for gene regulatory networks (GRNs) that could divide the petal surface into proximal and distal domains to create a bullseye, a very common type of petal pattern across the angiosperms. The evolved GRNs showed robust patterning dynamics and could generate a variety of bullseye proportions. We found that the evolution of bullseye patterns was often accompanied by the spontaneous emergence of a third cell type with a unique gene expression profile at the boundary between the proximal and distal regions. These bullseye boundary cells appeared in most simulations despite not being explicitly selected for, and we validated their presence experimentally in *Hibiscus trionum*, a model system whose flowers display a bullseye pattern. Although boundary cell types emerged spontaneously in our simulations, they evolved more often and were more important for pattern formation when gene expression was modelled as a noisy process. This suggests that GRNs producing this emergent cell type may support reproducible bullseye formation by buffering against developmental variability. Altogether, the results from our evolutionary simulations illuminate the early steps of petal pattern formation and demonstrate that novel cell types can arise spontaneously and repeatedly from selection on other cell types when developmental robustness is considered.

## Introduction

Flowering plants display a remarkable diversity of colourful and structural patterns on their petal surfaces, including spots, venations, bullseyes, and more complex, composite patterns. ^1^ These patterns play important roles in determining a flowering plant’s reproductive success, influencing both biotic interactions, like pollinator attraction and guidance, ^2–4^ and abiotic factors, such as protecting reproductive organs from harmful UV radiation or desiccation. ^5,6^ These roles are not mutually exclusive; in sunflowers, a larger UV-absorbing bullseye is associated with both reduced transpiration rates and increased pollinator attraction. ^6^ Thus, understanding how these patterns are formed during development and how they evolve is key to elucidating how morphological adaptation driven by different selection pressures originates.

While progress has been made in identifying the transcription factors (TFs) that directly drive epidermal cell differentiation in the different regions of the petal, ^7–10^ much less is known about the upstream regulatory events that divide the petal surface into distinct domains, allowing neighbouring cells to follow distinct developmental trajectories. ^1,11,12^ The spatial outlines of petal patterns are likely specified long before those motifs become visible on the petal surface, i.e., the petal is pre-patterned before structural genes are activated and cells acquire their characteristic features. ^4,13–16^ One recent hypothesis proposes that bullseye patterns, a very common type of petal pattern where the centre of the flower contrasts with its periphery, are initiated by an external signal that enters the developing petal primordium from its base, where it connects to the rest of the floral structure. ^4^

Previously, computational models of petal patterning have been developed to shed light on their developmental mechanisms. For example, Ding et al. ^17^ and Lu et al. ^18^ used reaction-diffusion models to simulate anthocyanin spot patterning in *Mimulus guttatus* and *Phalaenopsis* orchids, respectively, successfully incorporating and testing experimentally determined genetic interactions. Similarly, Ringham et al. ^19^ explored a wide range of floral patterns across different flower morphologies, driven by mechanisms including reaction-diffusion and positional information. These models demonstrate possible mechanisms by which fine-grained spots and stripes may appear in particular regions of the late-stage petal. However, it is still unclear how the discrete spatial domains of the petal are first established in emerging petal primordia.

To investigate the range of genetic mechanisms that could govern this early pre-patterning phase, we constructed a computational model of petal pattern evolution ^20–23^ and evolved GRNs that can transform a mobile signal into a stable bullseye pre-pattern across the surface of the petal primordium. To simulate the developmental dynamics of GRNs, we created a two-dimensional cell-based model of the petal epidermis where each cell can differentiate by expressing particular genes. We then selected for gene expression patterns resembling the distinct proximal and distal domains observed in petals with a bullseye pattern. ^2–6^ To further examine the expression profiles associated with the GRNs that successfully yield bullseye patterns, we developed a pipeline inspired by single-cell RNA sequencing (scRNA-seq) data analysis. This allowed us to investigate the evolution and development of different cell types across the petal surface.

We found that a third cell type evolved at the boundary between the proximal and distal regions in the majority of simulations. This cell type could develop via two distinct but not mutually exclusive expression profiles: (i) by preferentially expressing genes only in the boundary region, or (ii) partially overlapping gene expression domains, where the boundary forms in the non-overlapping region. We then demonstrated that such a bullseye boundary cell type is indeed present in the petal epidermis of *Hibiscus trionum*, a species recently established as a model system to investigate petal pattern development and evolution, and identified genes that are preferentially expressed in the bullseye boundary region during development in this species. To investigate the potential contribution of the boundary cell type, we performed targeted mutations in the GRNs evolved *in silico*, and found that the specification of the boundary cell type ensures the correct formation of the bullseye pattern. Finally, our simulations indicate that the evolution of the bullseye boundary cell type is influenced by the presence of molecular noise during gene transcription and translation; when such noise was absent, the boundary cell type persisted for fewer generations and was often not involved in establishing the proximo-distal pattern. These results suggest that boundary cells play a role in buffering developmental variability during bullseye pattern formation, revealing a novel aspect of petal patterning mechanisms.

## Results

We constructed a cell-based model of an early-stage petal epidermis (Figure 1A and Figure S1) to evolve GRNs capable of producing bullseye patterns where the proximal and distal regions eventually develop contrasting features. Genes in the GRN encode one of 12 unique proteins that can act cell-autonomously as TFs, diffusing TFs, or cell-cell communication factors (Figure 1B). Development is initialised with all gene expression set to zero, except for the genes coding for a diffusing patterning signal that initiates development. This signal is constitutively expressed in a subset of cells at the petal base (Figure 1A), representing the entry point of the signal into the petal epidermis. We model the signal as asymmetric along the medio-lateral axis because petals often show internal asymmetry (see Methods: Signalling condition). ^24,25^ To account for gene expression variability, we introduce noise in the molecular processes associated with the developmental model, such as gene transcription and translation.

**Figure 1.**
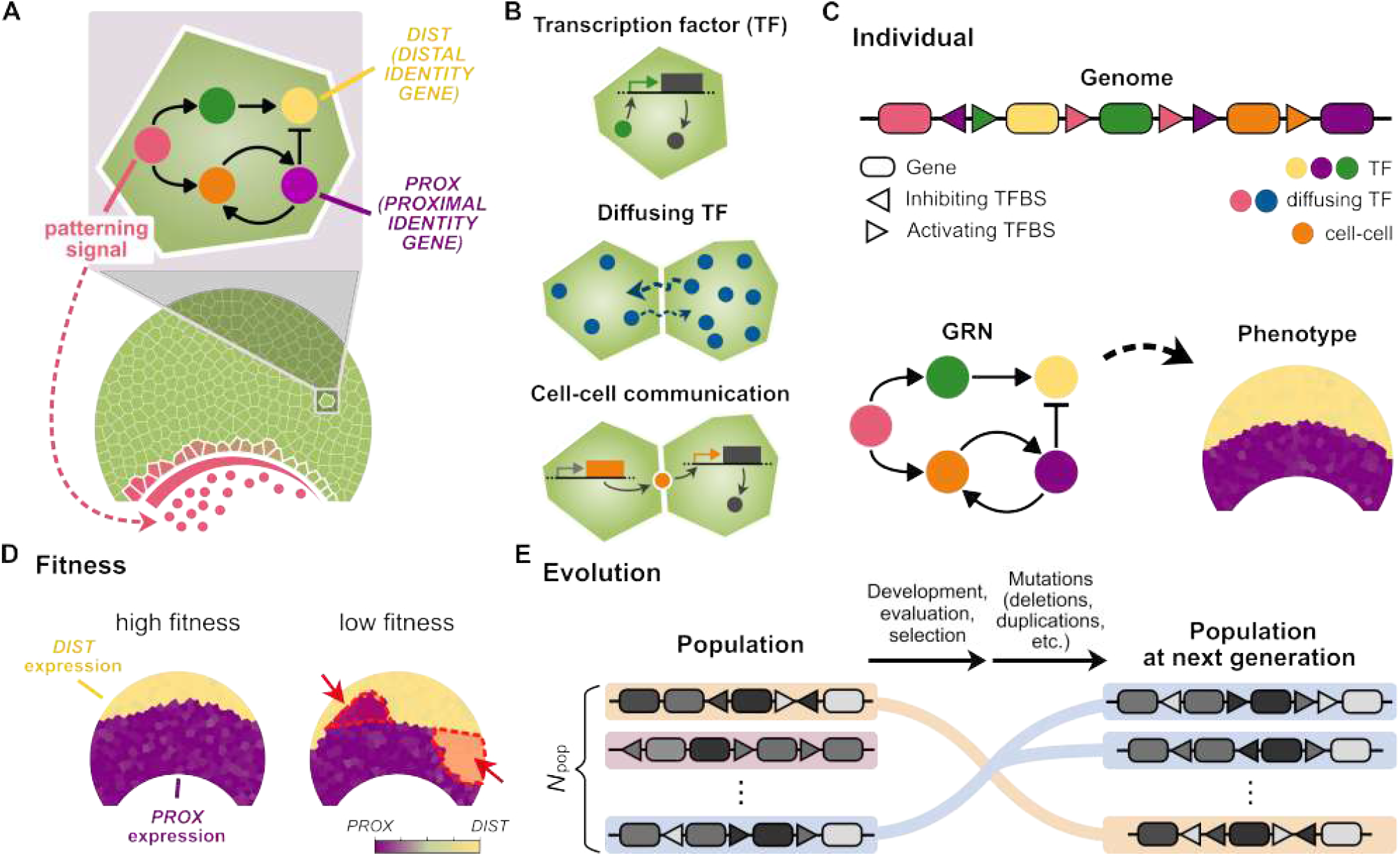
Computational model of petal patterning evolution and development. (**A**) A two-dimensional, cell-based model representing the petal epidermis. Gene regulation dynamics, governed by a GRN are simulated within each epidermal cell to drive the patterned expression of a *PROXIMAL IDENTITY GENE (PROX)* and a *DISTAL IDENTITY GENE (DIST)*. (**B**) The corresponding proteins can act cell-autonomously as TFs, or as diffusing TFs, or influence transcription in neighbouring cells only through cell-to-cell communication. (**C**) An individual’s genotype is represented by a linear genome comprising protein-coding elements (genes) and *cis*-regulatory elements (transcription factor binding sites (TFBSs)). This genome maps onto a GRN, where genes act as nodes and TFBS serve as edges, either activating or inhibiting interactions. The GRN dynamics are then simulated on the petal tissue to produce a phenotype. (**D**) Individuals are selected for a bullseye pattern, characterised by mutually exclusive expression domains of *PROX* and *DIST*. Examples of both high-fitness (left) and low-fitness (right) phenotypes are shown. In the low fitness example, unwanted *PROX* and *DIST* expression is marked in red. The proportion of the bullseye pattern is flexible and can evolve, ranging between 20% and 80% (proximal to total petal height). (**E**) Schematic of the evolutionary model used to evolve bullseye patterning mechanisms. In each generation, a population of *N*_pop_ = 1000 individuals undergoes development, with phenotypes evaluated and assigned fitness scores. Individuals selected for reproduction pass on their genome to their offspring (the new generation) with a probability of mutations. Mutations include duplication, deletion, innovation, and various changes in gene expression, dissociation rates, and regulatory weights (see Methods). These offspring will undergo development anew, and this cycle is repeated until the final generation *T*_gen_ = 30,000 is reached.

The developmental model determines the phenotype of individuals within an evolving population. Each individual in the population contains a beads-on-a-string-like genome encoding a GRN^26^ whose dynamics are simulated across the petal epidermis for a fixed amount of time, resulting in a phenotype (a gene expression pattern; Figure 1C). At the start of each evolutionary simulation, individuals in the population are randomly initialised with unique genomes: each genome contains one gene for each of the 12 gene types, but with randomly assigned regulatory interactions (GRN wiring) between them. After development, each resulting phenotype is assigned a fitness score based on how complete their bullseye pattern is: individuals obtain a high fitness score when their development results in a symmetric bullseye pattern, with the proximal region (centre of the flower bullseye) occupying between 20% and 80% of the petal height, where *PROXIMAL IDENTITY GENE (PROX)* is expressed in the proximal region and *DISTAL IDENTITY GENE (DIST)* in the distal region (Figure 1D). This is a proxy for the actual, more complex selection pressures on floral pattern arising from pollinator preference and abiotic factors. The fitness of the pattern is calculated by taking, for each cell, an average of its gene expression over a time window at the end of development, favouring stable expression patterns. Higher fitness scores result in a higher probability of reproduction, with offspring inheriting their parents’ genome with a probability of mutations (Figure 1E). Although the number of unique proteins remains fixed (12), the number of genes encoding them can change over evolution due to gene duplications and deletions.

We evolved 35 populations for 30,000 generations in independent evolutionary simulations, all of which successfully yielded GRNs producing robust bullseye patterns (Figure S2; see Figures S3 and S4 for all evolved GRNs). The distribution of maximum fitness achieved in the final generation across simulations is concentrated around a median fitness of approximately 60 out of a theoretical maximum of 100 (Figure 2A.i; see Figures S5 and S6 for evolutionary trajectories of population fitness and phenotypes, respectively). This deviation from the maximum primarily results from random fluctuations in gene expression and protein translation due to noise, as all evolved GRNs successfully create a bullseye pattern (Figure S2). We found that over a third of all simulations evolved a bullseye size of approximately 50% of the petal’s central height (Figure 2A.ii). This indicates a tendency for simulations to converge towards certain proportions more than others, possibly due to the interaction between the patterning signal distribution and the tissue geometry. Nevertheless, a broad range of bullseye dimensions did evolve across simulations (Figure 2A.ii), indicating the existence of GRN architectures that can give rise to variation in bullseye proportions.

**Figure 2.**
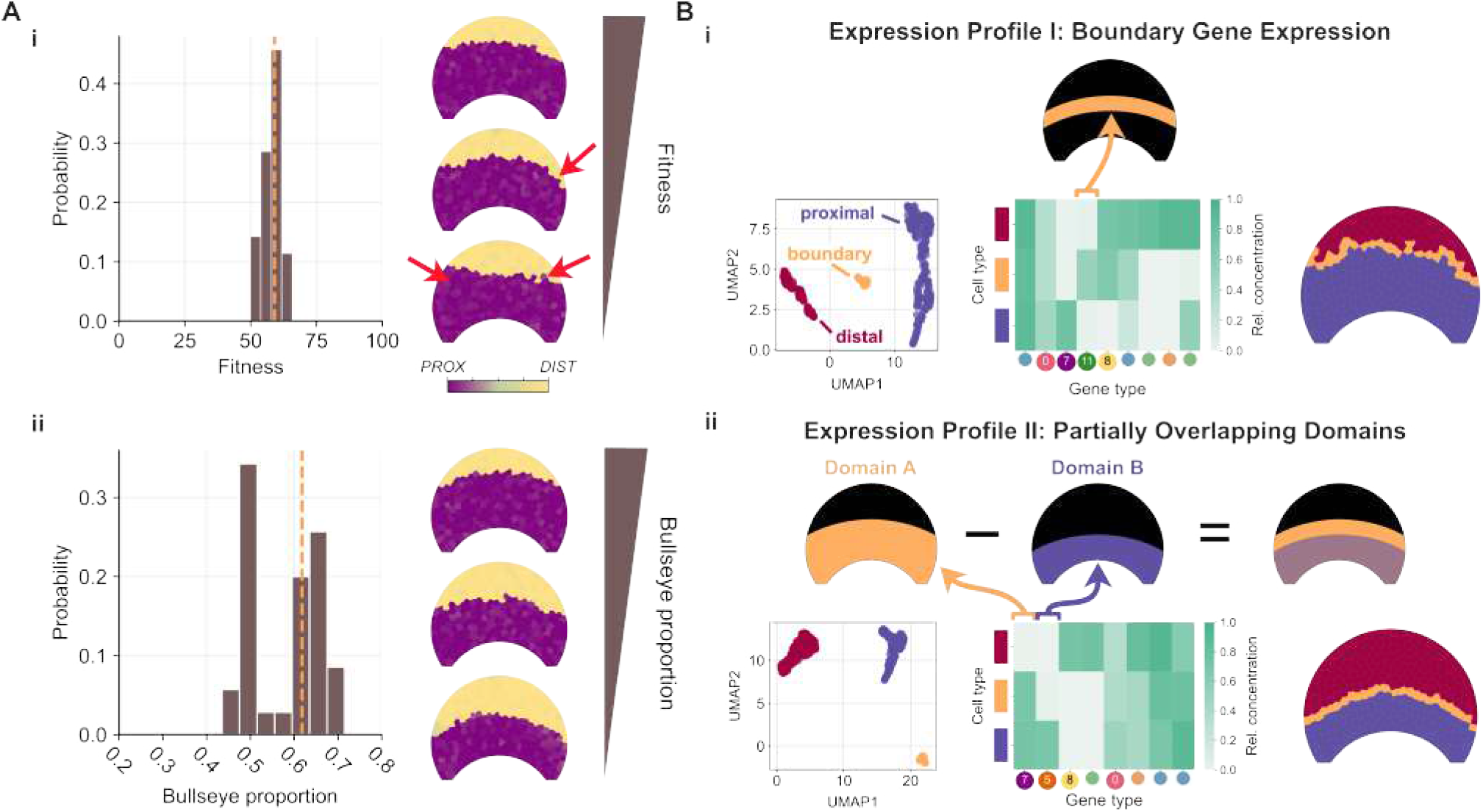
Populations readily evolve patterning mechanisms that spontaneously yield boundary cell types. (**A**) Overview of maximum fitness and bullseye proportions in evolved populations. (i) Distribution of maximum fitness achieved by individuals across all simulations. Examples of phenotypes from simulations with varying fitness are shown, with red arrows highlighting an incomplete bullseye pattern in the lowest-fitness simulation. Phenotypes are coloured by *PROX* and *DIST* expression levels (see Methods: *PROX* and *DIST* Visualisation). (ii) Overview of bullseye proportions in evolved populations, with example phenotypes from simulations exhibiting different bullseye proportions. Dotted vertical lines indicate medians. (**B**) Two distinct expression profiles drive the spontaneous evolution of boundary cell types. (i) Boundary emergence due to preferential expression of one or more genes in the boundary region (gene 11 in this example). From left to right: UMAP plot showing clustered cells, heatmap of relative protein concentration across the identified cell types, and spatial mapping of these cell types on the petal tissue. Concentrations are normalised relative to the maximum concentration of each gene type, and genes not differentially expressed are excluded from the heatmap. Node colours indicate molecular behaviour of each protein type (as defined in Figure 1B; see Methods). (ii) Boundary emergence from overlapping expression domains of uneven dimensions. In the example shown, gene 7 (*PROX*) is expressed in the larger bullseye domain (Domain A), while gene 5 is expressed in the smaller bullseye domain (Domain B). Figure S9 shows a complete overview of boundary cell types from all simulations in which they evolved.

### A third cell type evolves spontaneously at the bullseye boundary

To investigate how the genetic interactions in the evolved GRNs gave rise to bullseye patterning dynamics, we assessed the gene expression patterns of all genes involved in forming the bullseye pattern. We performed UMAP dimensionality reduction ^27^ on the evolved protein states, reducing the 12-dimensional protein space to two dimensions. Next, we performed HDBSCAN clustering ^28^ on this reduced space to identify different cell types across the petal epidermis (Figure S7; see Methods). For each simulation, we traced back the ancestral lineage of the final fittest individual and sampled 16 of its ancestors at evenly spaced generational intervals, performing this analysis on each sampled ancestor (see Figure S8 for a representative ancestral lineage). As expected, all lineages produced proximal and distal cells as our simulations explicitly selected for such cell identities. However, in 26 out of 35 lineages, we found individuals with a third boundary cell type, positioned between the two regions of the bullseye. In the remaining 9 lineages, successful bullseye patterning still evolved without this boundary cell type. Focusing on the majority of lineages in which this third boundary cell type arose, we analysed the gene expression profiles of the three identified cell types. This revealed two expression profiles by which the bullseye boundary cell type appears: (i) preferential or exclusive expression of certain gene(s) in the boundary region (expression profile I, Figure 2B.i), or (ii) genes displaying overlapping bullseye expression patterns with different proportions (expression profile II, Figure 2B.ii). Examples of developmental dynamics producing boundary cell types via these expression profiles are shown in Video S1 (expression profile I) and Video S2 (expression profile II).

Among the 26 simulations in which a bullseye boundary cell type evolved, expression profile II was the most common (15/26), followed by cases where both expression profiles co-occurred (8/26; Figure S14A). In contrast, expression profile I rarely evolved on its own (3/26). We hypothesise that expression profile I can emerge neutrally when expression profile II is already present to support the unique expression of a boundary gene in that narrow boundary region (Figure S10B).

### Cells at the *H. trionum* bullseye boundary express a unique set of genes during petal development

*Hibiscus trionum* (Malvaceae) has recently been developed as a model species to understand petal pattern development and its evolution. ^4,13,29^ This herbaceous species produces large flowers with a distinct bullseye pattern, defined by a purple proximal region contrasting with a cream-coloured distal domain (Figure 3A). Epidermal cells also vary in their shape and cuticle texture, ^4,29^ resulting in a pattern of distinct cell types along the petal proximo-distal axis (Figure 3A). Riglet et al. ^4^ found that the first morphological sign of cellular heterogeneity across the *H. trionum* petal epidermis appeared during the early pre-patterning phase (Stage 0 to 1), with the largest, most anisotropic cells emerging approximately one-third from the petal base (pre-patterning band). At this early stage, the epidermal cells have not yet begun producing pigment or acquired their characteristic shape and texture, but the position of the pre-patterning band coincides with the transition point between the future proximal, purple-pigmented cells and the distal, cream-pigmented cells of the final bullseye pattern. This suggests that those large anisotropic cells are destined to become the smooth and elongated tabular cells that characterise the bullseye boundary of mature petals (Figure 3A). ^4^

**Figure 3.**
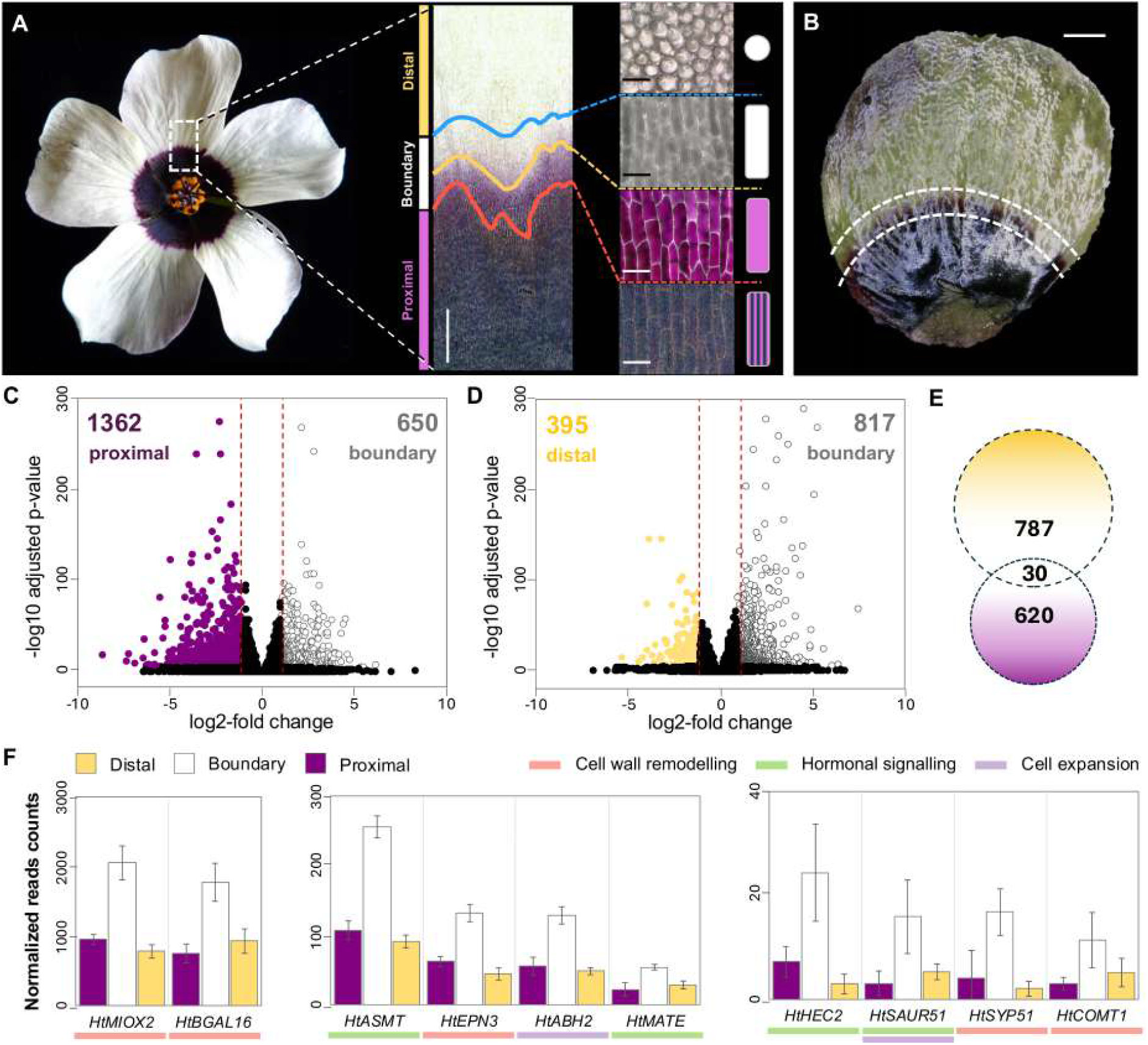
Comparative transcriptomics identifies genes preferentially expressed in the boundary region as bullseye emerges and supports the existence of a distinct boundary cell identity specified at early stages of petal development. (**A**) Close-up examination of the petal surface in mature flowers (stage 5) confirms the existence of a visible boundary region where epidermal cell features are distinct from the proximal and distal cells. Boundary cells are flat and elongated (tabular), but with a smooth surface. Cells in the lower boundary are anthocyanin-pigmented, while cells in the upper boundary are not. Symbols summarise cell types: striped purple rectangle = striated, pigmented, tabular cells; solid purple rectangle = smooth, pigmented, tabular cells; solid white rectangle = smooth, non-pigmented, tabular cells; solid white circle = smooth, non-pigmented, conical cells. Scale bars: central region close-up (left) = 500µm; individual cell types (right) = 50µm. (**B**) Dissected wild-type *H. trionum* Stage 2 petal showing the three regions used for RNA extraction and transcriptome analysis: distal (above top dotted line), boundary (between dotted lines), and proximal (below bottom dotted line). As the two morphologically distinct boundary cell types visible at maturity (A) have not yet differentiated at this stage, the boundary is treated as a single region. Scale bar: 1mm. (**C, D**) Volcano plots of genes from the stage 2 petal transcriptome analysis. Genes preferentially expressed (*>*2x) in the proximal, distal, or boundary region are shown as purple, beige, and white dots, respectively. Red lines indicate log2-fold thresholds; black dots indicate genes not differentially expressed. (**E**) Venn diagram depicting the genes preferentially expressed in the boundary compared to both proximal and distal regions. The number in the intersection indicates genes enriched in the boundary compared to both proximal and distal regions. The numbers within each non-overlapping portion of the circles indicate genes enriched in the boundary relative to only one region (proximal or distal), minus those shared in the intersection. (**F**) Expression levels of selected boundary genes involved in cell wall remodelling, hormone signalling, and cell expansion. Transcripts per million (TPM) values are shown; error bars = ±SD with *n* = 5 independent biological replicates. Subpanels are grouped by expression level: high (left), medium (centre), and low (right). *MIOX2* = *MYO-INOSITOL OXYGENASE 2, BGAL16* = Beta-galactosidase16, *ASMT* = *N-ACETYLSEROTONIN O-METHYLTRANSFERASE, EPN3* = *EPSIN3, ABH2* = abscisic acid 8’-hydroxylase, *MATE* = MATE transporter, *HEC2* = *HECATE2, SAUR51* = *SMALL AUXIN UP-REGULATED 51, SYP51* = *SYNTAXIN 51, COMT1* = *CATECHOL-O-METHYLTRANSFERASE 1*.

To test whether cells from the pre-patterning band correspond to the bullseye boundary cell type predicted by our simulations, we dissected Stage 2 petal primordia (the first stage where the pigmented bullseye becomes visible) and used RNA-seq to compare gene expression between the proximal, boundary, and distal petal regions (Figure 3B). Consistent with our model predictions, we found 650 and 817 genes expressed at least twice as much in the boundary region compared to the proximal or distal regions, respectively (Figure 3C,D; Tables S3 and S4). Most of these genes behaved like *DIST* (gene 8) from Figure 2B.i and *PROX* (gene 7) from Figure 2B.ii (or gene 7 from Figure 4B.i), while 30 showed boundary-specific expression (Figure 3E) resembling the profile predicted for gene 11 in Figure 2B.i. Thus, although only two epidermal cell types (anthocyanin-pigmented vs. anthocyanin-free) are readily distinguishable at Stage 2, a third boundary cell type is already present, characterised by unique gene expression patterns as predicted by the model.

**Figure 4.**
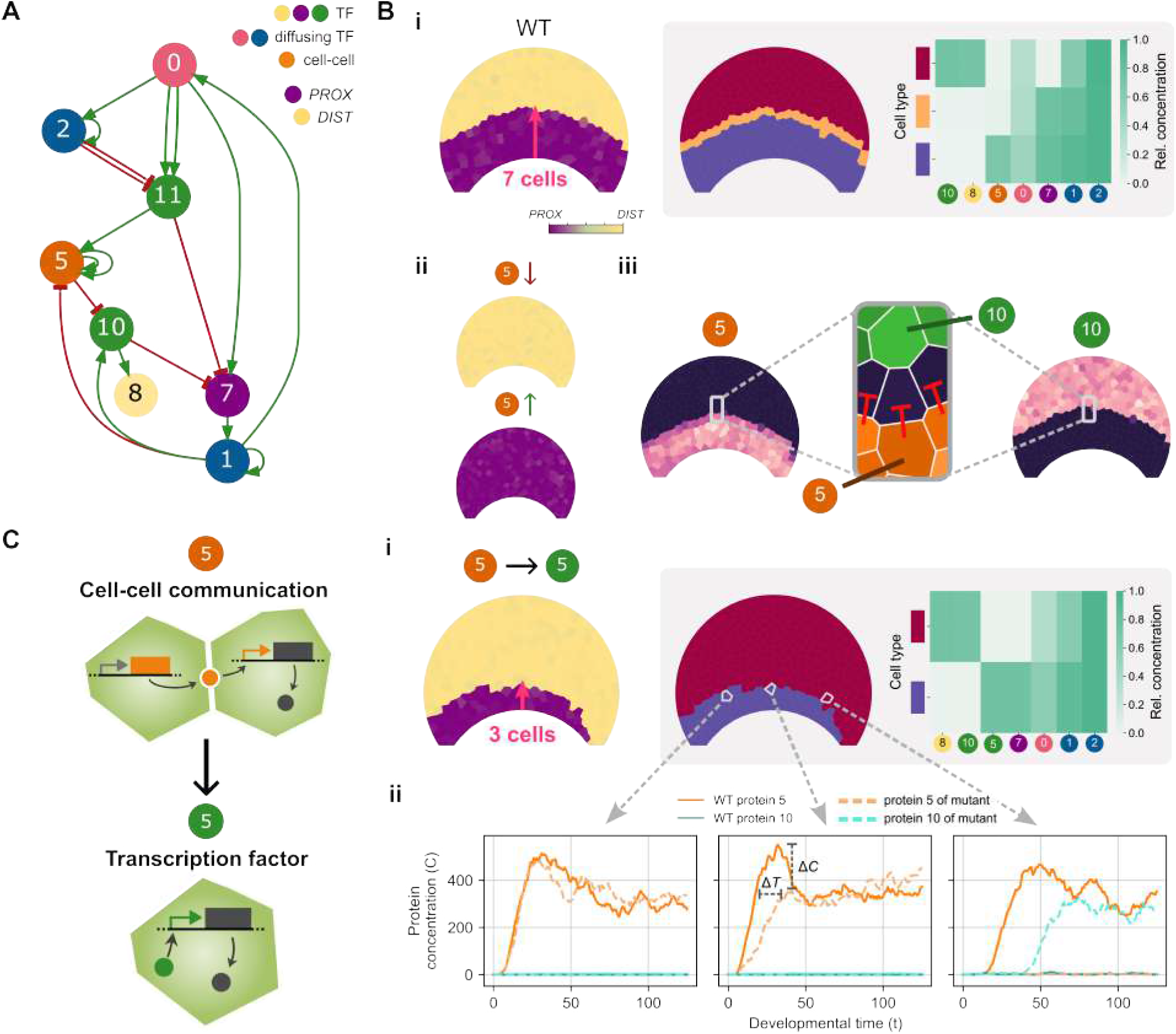
Boundary cell type plays a key role in correct bullseye patterning. (**A**) Pruned GRN of an individual from a representative simulation at generation 30,000. Vertex labels indicate gene types. Multiple arrows between nodes reflect multiple TFBSs, resulting in a stronger regulatory effect on transcriptional activity (see Methods). (**B**) Functional analysis of gene 5 in the GRN. (i) Wild-type phenotype showing normal bullseye pattern formation. (ii) Phenotypic effects of gene 5 knockout and overexpression, both leading to bullseye loss. (iii) Gene 5 restricts gene 10 expression to the distal region via cell-cell signalling, establishing a boundary inhibition zone essential for pattern formation. (**C**) Mutagenesis experiment converting gene 5 from a cell-cell communication gene, which influences neighbouring cells’ transcription, to one coding for a transcription factor (TF) acting cell-autonomously, only influencing transcription in cells in which it is expressed. (i) Resulting mutant phenotype showing loss of boundary cell type and bullseye symmetry. (ii) Temporal gene expression of genes 5 and 10 in the wild type and mutant, shown for three cells within the proximal bullseye region along the mediolateral axis. In the mutant, expression of gene 5 is both delayed (Δ*T*), reduced in magnitude (Δ*C*), and eventually lost with increasing distance from the signal origin.

Among the 30 genes whose expression peaks in the boundary domain of developing petal primordia (Stage 2), 7 encode hypothetical proteins, making their potential roles in bullseye formation difficult to predict. However, over half of the remaining genes (13 out of 23) encode proteins involved in cell wall remodelling, cell expansion and/or hormonal signalling in other species (Figure 3F; Table S5). For example, *MYO-INOSITOL OXYGENASE* catalyses the synthesis of glucuronic acid, a key precursor for cell wall polysaccharides, ^30^ while Beta-galactosidase16 participates in cell wall biogenesis and remodelling during organ growth. ^31–33^ *EPSIN3* participates in clathrin-mediated endocytosis, a process vital for wall deposition in apical growing plant cells; ^34^ SYNTAXINs play crucial roles during exocytosis, thus contributing to cytokinesis and the formation and maintenance of cell wall structure, and *CATECHOL-O-METHYLTRANSFERASE* contributes to the synthesis of lignin. ^35,36^ *ABH2* contributes to ABA metabolism, a hormone known to regulate cell elongation. ^37^ SAURs are regulators of (polar) cell expansion and mediate auxin-induced acid growth by promoting phosphorylation of plasma membrane H^+^-ATPase. ^38,39^ MATE proteins are also involved in the transport of phytohormones like auxin and abscisic acid. ^40^ Given the characteristics of the bullseye boundary cells, these processes are all likely to contribute to their initial specification during the pre-patterning phase and/or to their subsequent elaboration during petal development (Stage 1 to 5). Taken together, our results indicate that the third cell type predicted to frequently evolve by our *in silico* simulations is biologically relevant, as such cells are present in *H. trionum*, a model species for which the pre-patterning phase has been characterised.

### Contribution of evolved boundary cell type to *in silico* bullseye formation

Next, we investigated whether the very early boundary cells that evolved in our simulations are simply a frequent by-product of bullseye-producing GRNs, or whether this boundary cell type is important for petal pattern formation. We analysed the pruned GRN of a fit individual at generation 30,000 from a representative simulation in which a boundary cell type evolved via expression profile II (Figure 4A; a similar analysis is carried out on a non-boundary forming GRN in Figure S11). In this pruned network, genes and interactions with minimal impact on fitness were removed (see Methods for details and Figure S12 for examples of the pruning process). We identified gene 5, a cell-cell communication gene, as a key regulator of bullseye boundary cell type formation in this simulation. This gene is expressed in the proximal petal region, following a bullseye pattern one cell smaller in height than the bullseye pattern defined by the expression domain of *PROX* (Figure 4B.i). Disrupting the function of gene 5 by either knockout or overexpressing it constitutively across the petal led to uniform expression of *DIST* and *PROX*, respectively (Figure 4B.ii). In both cases, the bullseye pattern was lost, indicating that gene 5 is required for correct cell differentiation along the proximo-distal petal axis.

Gene 5 is a repressor of gene 10, which is a regulator of distal identity that is preferentially expressed in the distal region, activating *DIST* and inhibiting *PROX* expression (Figure 4A). By restricting the expression of gene 10 to the distal domain and establishing a boundary inhibition zone through its non-cell-autonomous behaviour (Figure 4B.iii), gene 5 acts as a promoter of proximal identity. To further investigate its contribution to bullseye formation, we generated an *in silico* mutant by modifying gene 5 to function as a TF acting cell-autonomously instead of a cell-cell communication gene (Figure 4C). This mutant produced a smaller and asymmetric bullseye pattern, lacking the bullseye boundary cell type (Figure 4C.i). Decreasing the spatial range of gene 5’s regulatory influence by turning it into a TF resulted in a delay in its inhibition of gene 10 and reduced its self-activation range, explaining the smaller bullseye. In this mutant, expression of gene 5 is progressively delayed in cells located further from the origin of the patterning signal, and is ultimately absent on the right side of the proximal region of the bullseye (Figure 4C.ii). As a consequence, gene 10 becomes expressed in the right region, resulting in *DIST* identity instead of *PROX*, and leading to an asymmetric bullseye pattern.

These *in silico* experiments also indicate that the bullseye is specified through a precise timing mechanism. Gene 5 delays and spatially restricts the expression of gene 10, ensuring the symmetric development of the pattern. Without this extended inhibition, boundary cells are lost, and the bullseye becomes misshapen, underscoring that this GRN generates a bullseye pattern through the creation of a boundary cell type.

### Molecular noise increases evolutionary persistence of bullseye boundary cell types

To better understand the evolutionary dynamics of the bullseye boundary cell types, we measured the number of generations for which they persisted in each simulation (boundary persistence times (BPTs); Figure 5A.i), using persistence as a proxy for functional relevance: boundary cell types that contribute to bullseye patterning are expected to be conserved, whereas those that arise without a patterning role are more likely to disappear quickly through fitness-neutral mutations. We find that boundary cell types are most commonly short-lived, persisting for 500 generations or less. However, in several simulations, this cell type persists over long evolutionary times (BPTs longer than 10,000 generations; Figure 5A.ii). To test whether these long-lived boundary cell types serve a developmental role, we pruned the genomes of individuals in simulations in which boundary cell types evolved. After pruning, boundary cell types either remained (see Figure S12A) or were lost (Figure S12B). In simulations with long BPTs, we found that the boundary cell types remained. Since pruning retains only the genes and interactions essential for bullseye pattern formation, this suggests that in these cases, boundary cells are necessary to establish proximo-distal differentiation.

**Figure 5.**
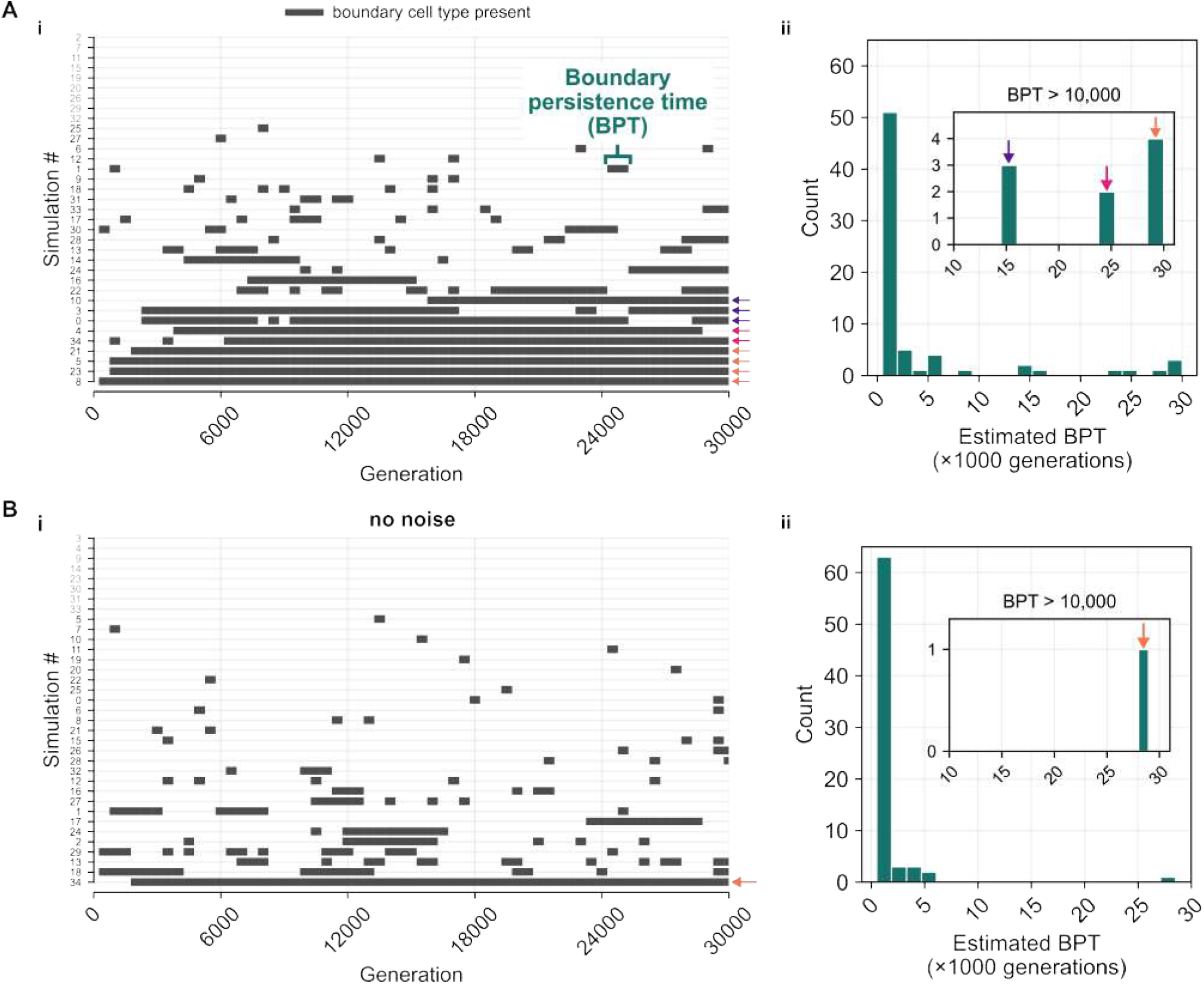
Boundary cell types persist longer during evolution in the presence of molecular noise. (**A**) Evolutionary dynamics of boundary cell types in simulations with molecular noise. (i) Estimated presence of boundary cell types across simulations over 30,000 generations. Boundary cell presence is assessed every 500 generations, introducing a margin of uncertainty of approximately 500 generations. Arrows highlight long BPTs: ∼30,000 generations (orange), ∼25,000 generations (magenta), and ∼15,000 generations (purple). The evolutionary trajectory of Simulation #34, including the progression of its phenotype and cell types, is shown in Figure S8. (ii) Distribution of BPTs across simulations. A single simulation may contribute multiple BPTs if boundary cell types reappear throughout evolution, or none if no boundary evolved. The inset provides a zoomed-in histogram of long BPTs. (**B**) Evolutionary dynamics of boundary cell types in simulations without molecular noise. (i) Presence of boundary cell types across simulations. The evolutionary gain and loss of the highly transient boundary cell type in Simulation #13 is shown in Figure S16, where we demonstrate that, in this particular simulation, boundary cells arise through multiple independent and diverse mutations rather than repeated rediscovery of a single change, explaining their frequent appearance. (ii) Distribution of BPTs across simulations.

Recent findings in *H. trionum* indicate bullseye boundaries can be specified very early in petal development. ^4^ To this end, we hypothesised that boundary cells play a role in making the patterning process more robust by separating the two main petal regions early on. Here, we tested this by performing 35 additional evolutionary simulations without molecular noise, and compared the frequency and distribution of BPTs (Figure 5B; see Methods). Bullseye boundary cell types evolved with similar frequencies in both conditions (27/35 simulations without noise vs. 26/35 simulations with noise). However, in most simulations without noise, boundary cells evolved more transiently, appearing and disappearing repeatedly through generations (Figure 5B.i). Additionally, in the absence of noise, we found a notable decrease in the frequency of long BPTs (Figure 5B.ii), suggesting that boundary cell types arise primarily through neutral changes rather than conferring a developmental function or selective advantage. Finally, when we pruned networks that produced a bullseye boundary cell type in simulations without noise, the boundary cell types mostly disappeared. This suggests that the boundary cell type was a neutral feature of these GRNs and were not necessary to produce a bullseye pattern. In contrast, a greater proportion of boundary cell types persisted after pruning in simulations with noise (Figure S13), consistent with the fact that simulations with noise also have longer BPTs. Together, these findings support the idea that GRNs producing boundary cells are more likely to be maintained under molecular noise, suggesting those GRNs have an increased capacity to produce robust outcomes, buffering against developmental variability.

Next, we examined whether the presence of molecular noise is associated with differences in GRN motif frequency. We find that *DIST* inhibiting *PROX* is more common than the other way around in both simulations with and without noise (Figure S15A). Interestingly, mutual inhibition between *PROX* and *DIST* is more common in the absence of noise (Figure S15A). Additionally, positive autoregulation is more abundant under noisy conditions (Figure S15B), opposing the common view that positive feedback increases response times and expression noise. ^41^ On the other hand, negative autoregulation is more abundant without noise, which is contrary to previous findings that negative autoregulation reduces gene expression noise. ^42–44^

We also compared how often each of the two expression profiles involved in bullseye boundary cell formation (Figure 2B) evolved in simulations with and without noise. While expression profile II is most prevalent in the presence of noise (five times more frequent than expression profile I in simulations yielding the boundary cell type; Figure S14A), expression profiles I and II evolved with the same frequency when noise is absent (Figure S14B). Hence, there appears to be no inherent bias towards one expression profile over the other when developmental variability caused by molecular noise is missing. Furthermore, the co-occurrence of both expression profiles was less common than in noisy simulations (Figure S14A,B). These results support our hypothesis that expression profile I by itself is unstable in the presence of molecular noise.

## Discussion

Bullseye patterns, a petal motif widespread across angiosperms, are thought to be specified by early developmental pre-patterns that direct the differentiation of distinct cell types. ^10,12,17,45^ However, the genetic mechanisms that specify these pre-patterns remain to be understood. ^46^ To start investigating how GRNs establish such early gene expression domains, we developed a cell-based model of the petal epidermis and *in silico* evolved GRNs capable of robustly generating bullseye expression domains. We analysed the successful GRNs with an approach inspired by scRNA-seq data analysis to identify the different cell types present. While our selection criterion specified only two cell types arranged in a bullseye pattern, a third cell type spontaneously emerged at the bullseye boundary in most simulations. We demonstrated that such a bullseye boundary cell type exists in the region that separates proximal and distal domains in *H. trionum* petals: we identified a set of genes preferentially expressed in the bullseye boundary domain in early stages of petal development (Stage 2), long before those cells exhibit the unique combination of structural features (flat tabular shape with a smooth cuticle) that distinguish them in the mature flower. Finally, by comparing simulations with and without molecular noise in the developmental model, we found that boundary cell types are likely not just a neutral side-product of bullseye formation, but instead their establishment could contribute to developmental robustness by buffering against variability in gene expression and other noisy molecular processes. The recent findings from Riglet et al., ^4^ showing that the *H. trionum* petal is pre-patterned with future bullseye boundary cells differentiating first, is also in agreement with our model and further emphasises the importance of such cells to pattern formation.

Using a transcriptomic approach, we identified 30 genes preferentially expressed in the bullseye boundary region at Stage 2 of petal development in *H. trionum*. These genes fit the predictions of expression profile I and provide experimental support for the existence of a third, distinct boundary cell type, central to bullseye formation. Whether the 30 bullseye boundary genes identified at Stage 2 are involved in the establishment of the bullseye boundary during the pre-patterning phase (Stage 0 to 1, as defined in Riglet et al. ^4^) or only act later to control the differentiation of the boundary cells (smooth tabular cells seen in Stage 5) remains to be tested experimentally. However, the known functions of their homologs in other species (cell wall remodelling, hormone signalling, and cell expansion) are consistent with roles in both the emergence and subsequent differentiation of the bullseye boundary.

Boundary establishment plays an important role during plant morphogenesis. ^47–49^ Cells in boundary zones are characterised by distinct gene expression profiles and typically exhibit reduced proliferation and growth rates, often contributing to individualisation of emerging primordia or influencing organ shape. ^49^ The best-characterised boundary genes in plants belong to the NAC (*NAM, ATAF1/2*, and *CUC2*) transcription factor family. ^50^ Loss-of-function mutations of NAC genes cause fusion of adjacent organs ^51–56^ and leaflets, ^56,57^ highlighting an important role for boundary genes in maintaining proper organ and leaflet separation during development. We did not find canonical NAC genes preferentially expressed in the boundary at the Stage 2 petal. However, their potential involvement at earlier developmental stages cannot be excluded, especially since these boundary cells stop dividing early on, which is characteristic of NAC gene activity in organ boundary cells. ^51,52^ Our simulations further suggest that boundary cells may have additional roles in pattern formation. Specifically, we found that boundary cell types may play a role in establishing symmetric bullseye patterns and may buffer patterning processes against molecular noise. However, boundary cells do not seem to be strictly required for bullseye pattern formation since we also found GRNs that generate bullseyes without boundary cells. These results suggest that boundary zones may serve not only structural roles in separating organs during morphogenesis, but also directly contribute to patterning events, playing dynamic roles in ensuring the stability and precision of cell fate specification across a tissue.

Interestingly, a similar boundary pattern was recently reported in the leaves of *Mimulus verbenaceus*, where the patterned expression of three TFs along the proximo-distal leaf axis gave rise to a pigmentation stripe. ^58^ One of those TFs acts as a proximal identity gene: it is preferentially expressed in the base of emerging leaves and inhibits the activator of pigmentation. ^59^ Knockout of this gene transformed the leaf pigmentation stripe into a bullseye-like pattern. This suggests that a combination of boundary expression profiles I and II is at play in which the proximal identity gene’s smaller bullseye expression domain restricts the activity of the pigmentation activator, otherwise expressed across a larger bullseye region, to the boundary domain, generating the purple leaf stripe. We found that such a combination of expression profiles is more likely to evolve and be functional in pattern formation when molecular noise is present, suggesting that the boundary pattern in *M. verbenaceus* leaves could initially have emerged to provide developmental stability, later gaining the ability to regulate the expression of pigment biosynthesis genes. Given the shared evolutionary origin of petals and leaves, ^60–62^ and the comparable proximo-distal organisation observed in both the *M. verbenaceus* leaf pattern and the *H. trionum* petal bullseye, it is possible that boundary cell types are an ancient feature of proximo-distal patterning processes at work in flat lateral organs. Alternatively, as *M. verbenaceus* and *H. trionum* diverged early in angiosperm evolution, around 125 million years ago, ^63^ they could also have evolved boundary cells independently via replicated evolution, as we demonstrate that boundary cell types can readily and repeatedly evolve in bullseye patterning systems.

The bullseye pre-pattern we investigate here is likely established through a fundamentally different mechanism than the downstream anthocyanin patterning studied by Ding et al. ^17^ and Lu et al. ^18^ Those models describe how fine-grained spots and stripes emerge from a uniform initial state via reaction-diffusion mechanisms, whereas our work addresses the earlier step of how an asymmetric signal is converted into discrete spatial domains. Our results thus complement these studies by providing candidate mechanisms for the pre-patterning events that establishes the regional domains within which self-organised patterns are subsequently confined.

Other theoretical studies have identified several qualitatively different GRNs that can produce stripes, similar to the boundary cell identified here, either through an exhaustive search of small networks^64,65^ or through an evolutionary process. ^20,21,66–68^ These studies showed that there are several qualitatively distinct mechanisms that can produce stripes that differ in their robustness, and whose evolution depends on the presence and type of positional information available to cells in the tissue. Stripe-forming mechanisms identified by such explicit selection could therefore be considered a form of between-level novelty *sensu* Colizzi et al., ^69^ where the stripe itself is assumed, but the novelty lies in the process (GRN) that produces it. Here, and in our recent work on evolution of stem-cell niche patterning in plants, ^23^ we find that selection on a particular pattern can also result in novel, not explicitly selected-for cell types that either support the development of the trait under selection (a bullseye or stem cell niche pattern), or simply emerge as a neutral by-product. These novel cell types are a clear example of what Gould and Lewontin ^70^ termed spandrels: traits that arise as by-products of developmental architecture or constraint, rather than evolving as direct adaptations. Importantly, our study provides a plausible mechanism of how such features could come about in the case of petal patterning. Our results also highlight the apparent ease with which evolution can generate boundary cell types in bullseye patterning systems, complementing the earlier findings from Jiménez et al. ^65^ that stripe-forming mechanisms have bullseye (high-low) patterns in their mutational neighbourhood. Our work therefore provides additional support for the idea that the inherent structure and constraints of a developmental system may bias evolution towards specific novel patterns. ^71–73^

Finally, our work focused on GRNs of pattern formation in a static tissue; however, morphogenetic processes also influence the proportions of the bullseye pattern, with growth acting as a pattern modifier. ^4^ The final dimensions of the bullseye therefore result from the combined action of pre-patterning events and regulation of growth in different regions of the petal, with changes to either process potentially affecting bullseye size. ^12^ Tissue growth may also distort the initial pre-pattern, so additional regulatory feedback between patterning and growth processes may be required to maintain or refine the pattern during petal development. ^74^ Future work could explore this interplay of pattern formation and morphogenesis by incorporating cell growth and division into the developmental model and evolving patterning mechanisms in a dynamically growing and dividing tissue.

## Resource availability

### Lead contact

For further information and resources, please contact the lead contact, Steven Oud, at.

### Materials availability

This study did not generate new unique reagents.

### Data and code availability

- The coding and genomic sequences of all *H. trionum* genes mentioned in this study can be accessed via the genome of *H. trionum* available on GenBank (https://www.ncbi.nlm.nih.gov/datasets/genome/GCA_030270665.1/) using the HRI_ reference numbers provided in Table S5. The EDWA@ sequences from the reference transcriptome have been published as Supplementary Data 1 in Yeo et al. ^13^ Transcriptomic data associated with the comparative analysis of gene expression between proximal, boundary, and distal petal regions of *H. trionum* Stage 2 petal (accession numbers SAMN48289654 to SAMN48289683) have been deposited on GenBank SRA (https://www.ncbi.nlm.nih.gov/sra). The algorithms and codes used for RNA-seq data analysis and differential gene expression analysis have been published elsewhere, as indicated in the references provided in the Methods section.
- All original C++ source code and Python analysis scripts used in this work are publicly available at https://gitlab.developers.cam.ac.uk/slcu/teamrv/publications/oud_2026 and https://doi.org/10.5281/zenodo.18923037.

## Acknowledgements

We acknowledge support from the entire SLCU community including lab support, microscopy, horticulture, and facilities teams. We also thank Enrico Sandro Colizzi and the Sainsbury Laboratory Evolution Journal Club for the helpful discussions and insightful comments on the manuscript.

## Funding

This work was supported by grants from the Gatsby Charitable Foundation (G112566) and Isaac Newton Trust – School of Biological Sciences Seed Funding (RG58452) to RMAV; the Gatsby Charitable Foundation (RG92362 and G117782) and the Isaac Newton Trust/Wellcome Trust ISSF (RG89529) to EM.

## Author contributions

RMAV and EM conceived and designed the project. SO, PLJ, and RMAV designed the computational model. SO conducted the simulations and analysed simulation data; MMZ developed single-cell inspired analysis pipeline. MTSY and EM performed the experiments; JFW assembled the transcriptomes and performed the differential gene expression analyses between proximal/boundary and distal/boundary; EM analysed the data and identified boundary genes. SO, RMAV, and EM prepared figures and wrote the manuscript, with input from all other authors.

## Declaration of interests

The authors declare no competing interests.

## Methods

### Plant material and growing conditions

Wild-type *H. trionum* L. CUBG seeds (voucher CGE00046422) were sourced from the Cambridge University Botanic Gardens, Cambridge, UK. Seeds were imbibed in 90°C H_2_O for 10 min, germinated in the dark at 30°C for 48 h before being transferred to Levington High Nutrient M3 compost and grown under glasshouse conditions consisting of a 16 h light/8 h dark photoperiod at 25°C with a minimum radiance of 88 W.m^−2^.

### Tissue collection, RNA extraction and Illumina sequencing for *H. trionum* stage 2 petal transcriptome

Tissue was collected from five individual *H. trionum* CUBG plants to generate five biological replicates: for each biological replicate, Stage 2 (S2) petals (as defined in Moyroud et al. ^1^) were harvested and dissected to separate proximal, boundary, and distal regions. RNA extraction, RNA-seq library preparation and Illumina sequencing were performed as described in Yeo et al. ^2^ Raw reads (accession numbers SAMN48289654 to SAMN48289683) have been deposited on GenBank SRA (https://www.ncbi.nlm.nih.gov/sra).

### *H. trionum* CUBG stage 2 petal transcriptome assembly and differential gene expression analysis

RNA-seq reads were used to assess gene expression and compare transcript abundance between proximal, boundary and distal regions of Stage 2 (S2) petal primordia, as defined in Moyroud et al. ^1^ Transcriptome assembly and differential gene expression analysis were performed as described in Yeo et al. ^2^ Genes with log2-fold change < − 1 or > 1 and adjusted *p*-value < 0.05 were considered differentially expressed. Results of the differential gene expression are provided in Tables S3 to S5.

### Flower imaging

To produce Figure 3A, an open flower (Stage 5), as defined in Moyroud et al. ^1^, was collected in the morning and imaged on black velvet using a Panasonic DMC-FZ38 camera. To produce Figure 3B and Figure 3A, dissected petals from Stage 2 bud and open flower (Stage 5) were imaged with a Keyence microscope (VHX5000 model) fitted with a VH-Z20R or VH-Z100R lens.

### Developmental model

#### Cell and tissue representation

We based our developmental model on the petal of *Hibiscus trionum*, because its pre-patterning stage has been well characterised. ^3^ We represent the Stage 0b adaxial petal epidermis of *H. trionum* as a two-dimensional mesh of cells (Figure S1). We discretise the petal domain into *N*_*C*_ cells by generating a Voronoi tessellation from a set of cell-centre sites. The Voronoi diagram is computed using Fortune’s algorithm. ^4^ The resulting tessellation defines a planar polygonal mesh, which we interpret as a vertex-based tissue: vertices correspond to Voronoi vertices, edges to cell-cell interfaces, and each cell is represented by the polygon associated with its generating site (Figure S1D). This construction provides the geometric quantities (cell areas, edge lengths) required by the GRN model.

To account for slight variation in cell configurations in developing tissues due to noise, we generate *N*_morph_ distinct petal morphologies by adding a small amount of noise to each site’s position. During evolution, individuals are independently assigned a random morphology each generation, such that it will need to evolve mechanisms that are robust against variation in cell configuration.

#### Cell-cell interactions

Cells can affect the behaviour of neighbouring cells either by exchanging gene products through diffusion or by influencing a neighbouring cell’s gene expression directly through cell-cell communication genes. Neighbour relationships are defined by the Delaunay triangulation, i.e., the dual graph of the Voronoi tessellation used to generate the tissue. ^5^ In practice, each cell centre corresponds to a node in the Delaunay graph, and two cells are considered neighbours if their corresponding Voronoi polygons share a common edge. This defines the adjacency network used for intercellular interactions.

We assign a weight to each neighbouring cell pair proportional to the length of their shared Voronoi edge:

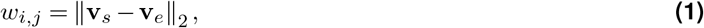

which can be thought of as the contact surface area of cell *i* and cell *j*’s cell walls. This weighted adjacency graph is then used to scale diffusion and cell-cell signalling terms in the GRN dynamics (Equations (6) and (8)).

#### Gene expression and regulation

Our developmental model is defined as a system of stochastic differential equations (SDEs) that describes the continuous change in mRNA and protein concentrations in discrete cells in the petal tissue. The dynamics of the system are given by a GRN, which is a directed graph 𝒢 = (*V, E*) that is determined by a genome. ^6^ A vertex *i* ∈ *V* (𝒢) represents a gene, which has a corresponding gene type *τ* (*i*) ∈ {0,…, 11} that determines the behaviour of the gene’s protein. The incoming and outgoing edges of a vertex then correspond to the gene’s incoming and outgoing regulatory interactions, respectively.

We model the forward flow from gene to mRNA to protein, with proteins, encoding transcription factors, regulating the mRNA transcription rates. In each cell, the regulatory dynamics are governed by the same GRN, whereas mRNA and protein concentrations can vary between cells. Each gene *i* in cell *j* has its mRNA concentration modelled by the chemical Langevin equation (CLE) ^7^:

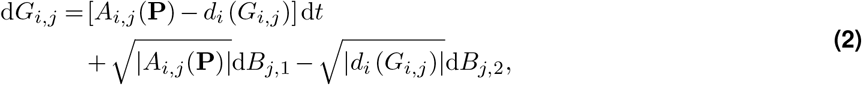

where *G*_*i,j*_ denotes the mRNA concentration of gene (vertex) *i* in cell *j* and *B* denotes a Wiener process. The promoter activity function *A*_*i,j*_ determines the transcription rate based on the transcriptional activity *T*_*i,j*_:

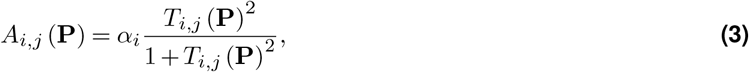

where *α*_*i*_ is the maximum mRNA transcription rate of gene *i*. The Hill function in Eq. (3) serves to map the unbounded total TF activity *T*_*i,j*_ at the promoter of gene *i* onto the interval [0, *α*_*i*_], ensuring that the transcription rate saturates smoothly to its maximum value *α*_*i*_, following Vroomans et al. ^8^

The decay function *d*_*i*_ describes the degradation of mRNA:

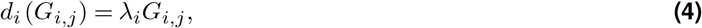

where *λ*_*i*_ is the mRNA decay rate of gene *i*.

The function *T*_*i,j*_ (**P**) describes the combined regulatory input of gene *i* by integrating the concentrations of the proteins for which it has one or more TFBSs:

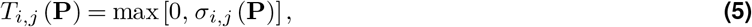

with the regulatory input *σ*_*i,j*_ a function of all regulatory sites in front of gene *i*:

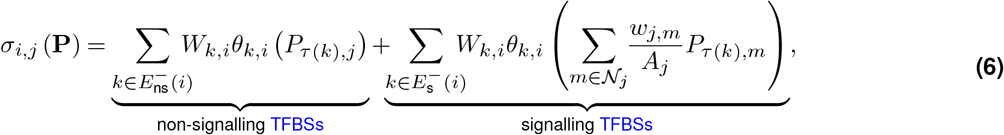

where

- *P*_*l,j*_ denotes the protein concentration of gene type *l* in cell *j*.
- 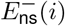 and 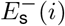 are the sets of non-signalling and signalling incoming edges of gene *i*, respectively.
- *W*_*k,i*_ is the edge weight which is −1 when an edge is repressive and 1 when an edge is activating.
- *θ*_*k,i*_(*x*) is the Hill function

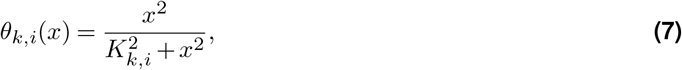

describing the activity of the regulatory element as a saturating function of the concentration of the TF with gene type *τ*(*k*) (*x* here), with *K*_*k,i*_ being the concentration of the TF at which it occupies half of the binding sites (*θ*_*k,i*_(*x*) = 0.5).

- 𝒩_*j*_ denotes the set of neighbouring cells of cell *j*, i.e., the vertices to which site **s**_*j*_ has an edge in the Delaunay triangulation.
- *w*_*j,n*_ is the interaction strength between cell *j* and *n* determined by their contact surface area (Eq. (1)).
- *A*_*j*_ is the area of cell *j* as calculated from the Voronoi tessellation by the shoelace formula.

We have separate terms in Eq. (5) for TFBSs with non-signalling and signalling gene types, as regulation by signalling gene types is dependent on the signalling gene’s protein concentrations in neighbouring cells. Altogether, Eq. (5) integrates all inhibiting (negative terms) and activating (positive terms) TFBSs of gene *i*, which determines the transcriptional activity of gene *i*. When the effect of inhibiting TFBSs is greater than the effect of activating TFBSs such that their sum is negative, the gene is considered to have no transcriptional activity, i.e., *T*_*i,j*_ (**P**) = 0.

Next, the dynamics of protein concentrations are given by the SDE

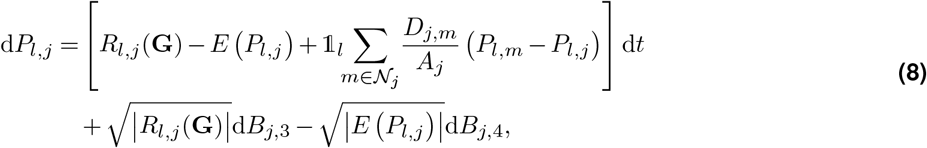

where *B* again denotes a Wiener process, and

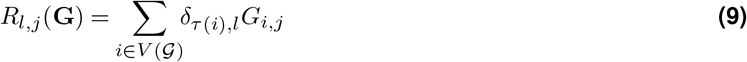

represents the translation of all mRNAs that have the same gene type *l*, i.e., all vertices in the GRN 𝒢 for which *τ* (*i*) = *l*. Here, *δ*_*τ* (*i*),*l*_ is the Kronecker delta which enforces the condition *τ*(*i*) = *l* (i.e., it is 1 if *τ*(*i*) = *l* and 0 otherwise). This acts as a filter to get all genes with gene type *l*. Next,

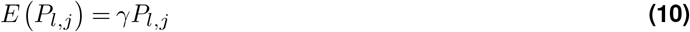

is protein decay where *γ* is the global protein decay rate, and *P*_*l,j*_ is the concentration of protein *l* in cell *j*.

Finally, in the diffusion term,

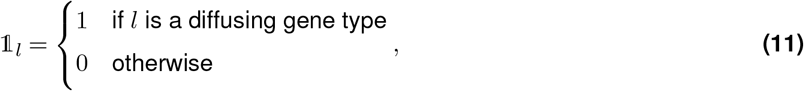

and *D*_*j,m*_ = *D*_0_ *w*_*j,m*_, where *D*_0_ is the base diffusion rate (Table S1), and *w*_*j,m*_ is the contact surface area of two cells *j* and *m* (Eq. (1)), such that cells with a larger contact surface area exchange proteins at a higher rate. As we have discrete cells whose interactions are described by an undirected graph (the Delaunay triangulation), diffusion is implemented as the discrete Laplace operator. Note that protein diffusion remains a deterministic process in this model.

These SDEs are solved using a stochastic equivalent of Ralston’s second-order Runge-Kutta method (for an example of the developmental dynamics, see Figures S1 and S2). ^9^ This method converges to the solution of the Stratonovich of our SDE and has a strong order of 1. ^10^ The diffusion matrix resulting from protein diffusion in Eq. (8) is solved using SuperLU, ^11^ a sparse direct solver that implements supernodal LU factorization. We integrate the system for a fixed developmental duration of *T*_*D*_ = 140 hours. Developmental time parameters (see Table S1) were chosen to approximately match the developmental window of *H. trionum* from stage 0 to stage 2, during which pre-patterning is established. ^3^

#### Deterministic developmental model

Deterministic ordinary differential equations (ODEs) are easily derived from the SDEs. For mRNA, we rewrite Eq. (2) as an ODE as follows:

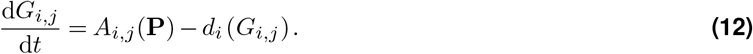

Similarly, for the proteins (Eq. (8)):

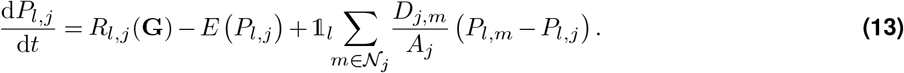

The resulting system of ODEs is numerically integrated using Ralston’s second-order Runge-Kutta method. ^9^

#### Gene types

Genes have a specific type, of which there are 12. Each gene type is identified by a unique number from 0 up to 11 (inclusive), and as a result of gene duplications, there may be several genes with the same gene type. However, there are always only 12 TFs, corresponding to the 12 unique gene types.

Individuals in the initial population are initialised with genomes that contain at least one gene of each gene type, and between one and three random TFBSs for each gene. Genes code for TFs that can affect the transcriptional activity of genes if they have a TFBS for the corresponding gene type. These TFs behave in different ways depending on their gene type:

- **Gene types 0, 1, and 2** code for diffusing TFs. These TFs are able to move to neighbouring cells through diffusion.
- **Gene types 3, 4, and 5** code for cell-cell communication TFs that influence the transcription of genes in neighbouring cells instead of the cell where they are expressed. This represents membrane receptor-mediated cell-cell communication, where a ligand (the TF) binds to a receptor protein bound to the membrane of neighbouring cells, which causes a signalling cascade that leads to a change in the target gene’s activity in the neighbouring cell.
- **Gene types 6-11** code for immobile TFs that simply affect the transcriptional activity of genes in the cells they are expressed in.

The number of gene types (and the number assigned to each behavioural category) was chosen to provide sufficient degrees of freedom while remaining computationally tractable. In most simulations, the pruned networks contain a subset of the 12 genes, suggesting sufficient degrees of freedom.

Special identity is assigned to certain genes. Gene type 0 represents the mobile signalling molecule which initiates the patterning of the petal. Gene type 7 represents *PROXIMAL IDENTITY GENE (PROX)*, i.e., a gene that is preferentially expressed in the proximal region of the *H. trionum* petal, and gene type 8 represents *DISTAL IDENTITY GENE (DIST)*, i.e., a gene preferentially expressed in the distal region of the petal.

#### Signalling condition

We chose to use an asymmetric signalling distribution as initial condition (i.e., the signal first enters the petal from one side of the organ base, generating a medio-lateral gradient) because petals often show internal asymmetry, ^12^ and initiation sites on floral meristems tend to be asymmetric, particularly in corollas with contorted aestivation that are widespread among flowering plants. ^13^ In addition, the behaviour of petal epidermal cells during the pre-patterning phase in *H. trionum* is also consistent with asymmetric signalling conditions. ^3^ Finally, from a theoretical viewpoint, while isotropic diffusion can be observed in living systems, asymmetric and non-uniform distribution of signalling molecules is much more likely in structured tissues due to cellular arrangement. ^14,15^

Gene type 0 is constitutively expressed in a small subset of cells near the petal base, representing the entry point of the patterning signal into the adaxial petal epidermis:

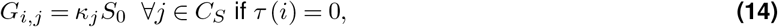

where *κ*_*j*_ ∈ [0, 1] is the intensity of the signal in cell *j, S*_0_ the baseline constitutive signal expression, and *C*_*S*_ the set of signal-producing cells at the base of the petal. The signal intensity decreases linearly with the *x* position of the cell:

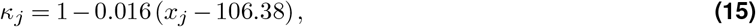

where *x*_*j*_ ∈ [0, 711.76] µm is the *x* position of the centroid (site) of cell *j*.

Note that the protein translation rate of a gene type is given by the sum of all the mRNA concentrations (Eq. (8)). Hence, having multiple copies of the signalling gene (gene type 0) increases the rate at which the signal enters the system (Eq. (14)).

#### Fitness function

Let *C* denote the set of all cells in the tissue, and *C*_*P*_ (*r*_*y*_) and *C*_*D*_ (*r*_*y*_) the subsets of cells belonging to the proximal and distal region of a bullseye with height *r*_*y*_, respectively. These subsets are disjoint (mutually exclusive) and their union is *C*:

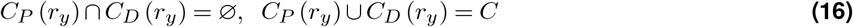

To select for a bullseye pattern, we want the PROX (gene type 7) protein at a certain target concentration *T*_*P*_ = 200 in the proximal cells, and the DIST (gene type 8) protein at target concentration *T*_*P*_ in the distal cells. Additionally, we want PROX proteins to have a concentration of 0 in the distal cells, and DIST proteins to have a concentration of 0 in the proximal cells.

Formally, we assign a score for how closely a protein concentration *x* matches the desired concentration in a cell *j* for a subset of cells *S* ∈ {*C*_*P*_, *C*_*D*_}:

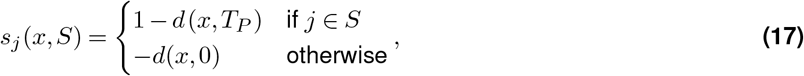

where

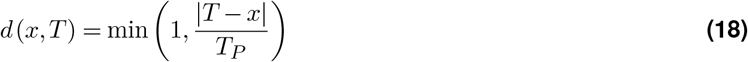

##### Algorithm 1

Bullseye Fitness Function *F* (*n, r*_*y*_)

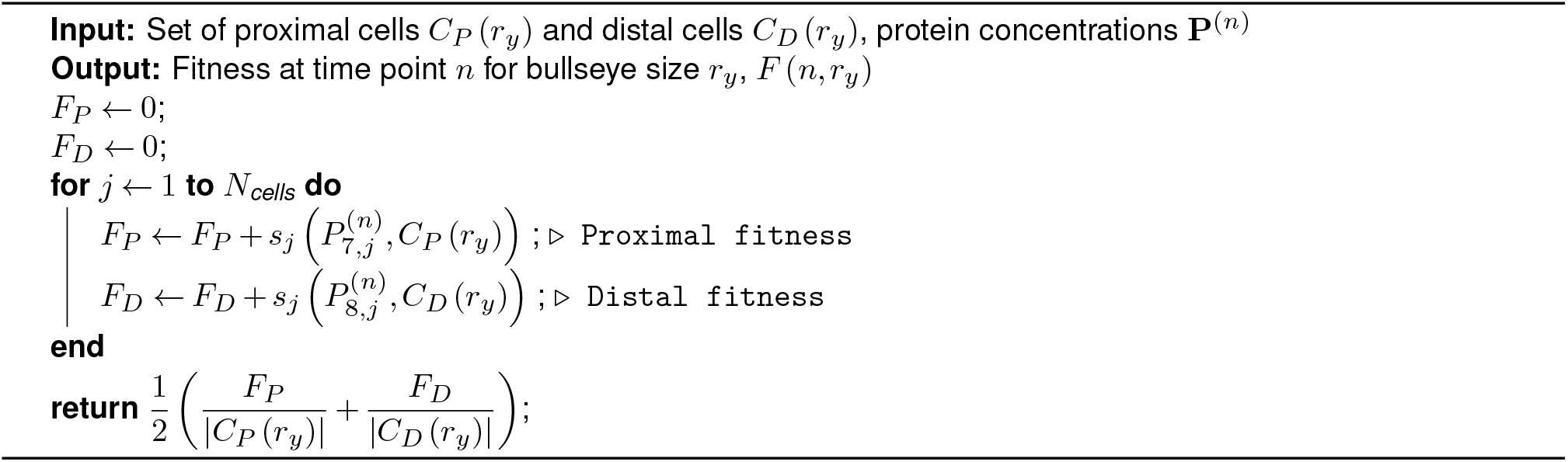

measures the normalised distance between the desired target concentration *T* and the actual concentration *x*. The fitness at a discrete time point *n* is then given by Algorithm 1. That is, an individual’s fitness is determined by the correct distribution (i.e., two mutually exclusive domains per Eq. (16)) of PROX and DIST proteins. An individual’s total fitness is the average of Algorithm 1 over a time window at the end of development (Table S1):

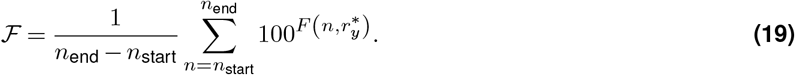

Note that as *F*(*n, r*_*y*_) ∈ (−∞, 1], the range of ℱ is (0, 100], i.e., the maximum achievable fitness is 100. While we do not explicitly select for gene expression stability, this averaging in Eq. (19) implies a penalty for unstable expression.

The proximal cells are defined as the cells inside of a horizontal ellipse with height *r*_*y*_ and width 1.02 · *r*_*y*_:

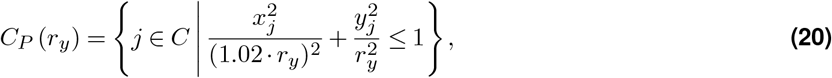

and the distal cells are the remaining cells outside the ellipse: *C*_*D*_ (*r*_*y*_) = *C \ C*_*P*_ (*r*_*y*_). The size of the bullseye pattern (i.e., the sets of proximal and distal cells) is determined by optimising the height *r*_*y*_ of the ellipse at developmental timestep *n* = *n*_start_ such that *F* (*n, r*_*y*_) is maximised. Local bounded scalar optimisation is performed using Brent’s algorithm ^16^ to determine the bullseye height:

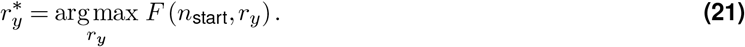

This optimised height 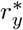 is then used to evaluate an individual’s fitness (Eq. (19)). Note that we restrict the range of possible values for 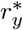 to between 20% and 80% of the petal’s height to ensure there is sufficient contrast between the proximal and distal domains, maintaining the appearance of a bullseye pattern.

We note that the choice of fitness function can influence simulation outcomes. We previously ran simulations with a fixed rather than dynamic bullseye size, and boundary cell types still evolved, suggesting our findings are robust to this variation. More substantial differences would be expected if selecting for abstract ecological criteria such as pollinator conspicuousness rather than a explicit spatial pattern; however, translating such criteria into a quantitative fitness function is a non-trivial challenge and outside the scope of this study.

#### Genome pruning

By representing the genotype as an explicit genome, the genetic architecture is flexible, where gene and TFBS duplications and deletions can create variation in genome size. Additionally, when a gene is duplicated, its upstream TFBSs are duplicated as well, allowing duplicated genes to change their upstream TFBSs, facilitating sub- and neofunctionalisation of duplicated genes. That is, evolution can tinker with the activity of the duplicated gene without affecting the original gene’s functionality. This open-endedness gives the evolutionary model more degrees of freedom, which is suggested to improve evolvability ^17^ but also generally leads to an increase in the size and complexity of evolved GRNs. To this end, we prune the evolved genomes, removing genes and TFBSs that have little to no effect on an individual’s fitness. The GRNs resulting from such pruning are referred to as pruned GRNs.

The pruning process is as follows. ^8,18,19^ We remove each gene and TFBS from the individual’s genome one at a time and evaluate the effect of this removal on fitness by repeating development 20 times and averaging the fitness outcomes. If the average fitness change is less than 3% of the individual’s fitness, the prune is accepted, and the gene or TFBS is removed from the genome. This process continues until no further deletions meet the 3% threshold. A depiction of this pruning process is shown on two representative networks from our simulations in Figure S12.

### Evolutionary model

The evolutionary model consists of a population of *N*_pop_ individuals, each of whom has a linear genome that contains transcription factor (TF)-coding genes as well as transcription factor binding sites (TFBSs) upstream of these genes that determine how their expression is regulated. ^6^ Individuals are initialised with a genome containing one copy of each gene type, and each gene has on average two TFBSs with random gene types and uniformly random activation or inhibition. The population is evolved for *T*_gen_ generations. At each generation, individuals undergo development, evaluation, and selection based on their fitness (i.e., how well they produce a bullseye pattern). Individuals are selected to reproduce, sometimes multiple times, and the offspring inherit the parental genome on which random mutations occur. That is, the next generation’s population is generated by randomly sampling parents with replacement, and thus a single parent may create multiple offspring (but with distinct random mutations).

#### Selection

Every generation, individuals undergo development in which the genotype is mapped into its corresponding phenotype. Based on how closely the phenotype of an individual *i* ∈ {1,…, *N*_pop_} resembles a bullseye pattern, they are assigned a fitness score ℱ_*i*_ (Eq. (19)). The new population is chosen by taking *N*_pop_ random individuals from the population, where the probability of an individual being selected is given by

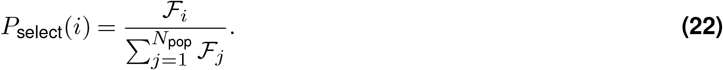

That is, the population size remains constant, and the probability of an individual being selected is proportional to their fitness. Each individual is sampled independently, and the same individual can be selected multiple times.

#### Mutations

After selection, each individual in the selected population can undergo multiple mutations. The following mutations can occur:

- **Gene duplication**. Each gene has a probability of *P*_genedup_ to duplicate itself and its TFBSs into a random spot in the genome.
- **Gene deletion**. Each gene has a probability of *P*_genedel_ to have itself and its TFBSs deleted. Note that at least one copy of each gene type is always maintained over evolution.
- **Gene maximum transcription rate change**. Each gene *i* has a probability of *P*_genetransc_ to change its maximum transcription rate *α*_*i*_. If the mutation occurs, the new maximum transcription rate becomes 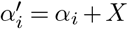, where *X* ∼ 𝒩 (0, *σ*_*α*_) is a random variable sampled from a normal distribution with mean 0 and standard deviation *σ*_*α*_.
- **Gene decay rate change**: Each gene *i* has a probability of *P*_genedecay_ to change its mRNA decay rate *λ*_*i*_. If the mutation occurs, the new mRNA decay rate becomes 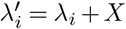 where *X* ∼ 𝒩 (0, *σ*_*λ*_).
- **TFBS duplication**. Each TFBS has a probability of *P*_TFBSDup_ to copy itself in front of a random gene.
- **TFBS deletion**. Each TFBS has a probability of *P*_TFBSDel_ to be deleted.
- **TFBS innovation**. There is a *P*_TFBSnew_ probability of inserting a TFBS with a random type and weight in front of a random gene.
- **TFBS dissociation constant change**. Each TFBS has a probability of *P*_TFBSChange_ to change its dissociation constant *K*_*k,i*_. If the mutation occurs, the new dissociation constant becomes 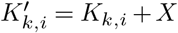 where *X* ∼ 𝒩 (0, *σ*_*K*_).
- **TFBS type change**. Each TFBS has a probability of *P*_TFBStype_ to change its gene type. That is, the identity of the source gene type from which the regulatory interaction originates in the network (i.e., the “from” node of the edge).
- **TFBS weight change**. Each TFBS has a probability of *P*_TFBSinv_ to invert its weight *W*_*k,i*_, i.e., go from acting as an activator to an inhibitor and vice versa. If the mutation occurs, the new weight becomes 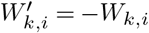.

Evolutionary parameters (see Table S1 for a comprehensive list) were based on previous studies using this modelling framework ^8,18,19^ and were adjusted during an initial exploratory phase to ensure a good balance between the strength of selection and speed of evolution.

### Cell Types and Dimensionality Reduction and Clustering

In this study, cell types are defined based on gene expression profiles. While the concept of cell identity remains an active topic of discussion, ^20^ we adopt a pragmatic definition in which distinct cell types correspond to qualitatively distinct expression states across the tissue. To identify such states, we use an automated dimensionality reduction and clustering procedure as a first-pass classification, followed by manual verification. This additional inspection ensures that identified cell types reflect genuinely distinct expression profiles rather than arbitrary partitions of smooth spatial gradients.

The dimensionality reduction and clustering procedure is as follows. The Uniform Manifold Approximation and Projection (UMAP) algorithm was used for dimensionality reduction using the UMAP Python library. ^21^ First, protein concentrations across the tissue were filtered to exclude proteins with low relevance. Specifically, proteins with a total concentration below 10 across the entire tissue or a standard deviation under 5 were removed, as these indicate low overall levels and uniform patterns, respectively. Next, each protein concentration was normalised to its highest expression level across the tissue, giving relative protein concentrations that range between 0 and 1. Finally, UMAP dimensionality reduction was applied to the normalised data. Two distinct UMAP embeddings were used: one for clustering and one for visualisation. The clusterable embedding uses the default UMAP parameters with n_neighbors = 30 and min_dist = 0.0, whereas the visualisation embedding uses n_neighbors = 30 and min_dist = 0.1. The same random state 555 was used for all UMAP embeddings to ensure reproducibility. Subsequent clustering was performed using the HDBSCAN Python library ^22^ to obtain cell types, which are visualised in a heatmap and ordered based on cell type and gene type using hierarchical clustering. HDBSCAN parameters used were min_samples = 15 and min_cluster_size = 20.

### *PROX* and *DIST* Visualisation

The colour of each cell is visualised by alpha-compositing two layers: *PROX* concentration (*c*_TF_ = purple) and *DIST* concentration (*c*_TF_ = cream) over a base green colour (*c*_base_). Each TF is represented by a colourmap with a transparent-to-opaque gradient, such that the RGBA value at concentration *x* is given by *f*(*x*) = (*r, g, b, α*(*x*)). The final displayed colour is computed as:

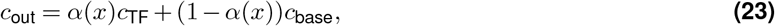

where *α*(*x*) ∈ [0, 1]. Both layers are computed independently and then composited over the base colour, such that cells expressing only the proximal factor appear purple, cells expressing only the distal factor appear cream, and non-expressing cells retain the base colour.

## Supplementary Information

**Figure S1.**
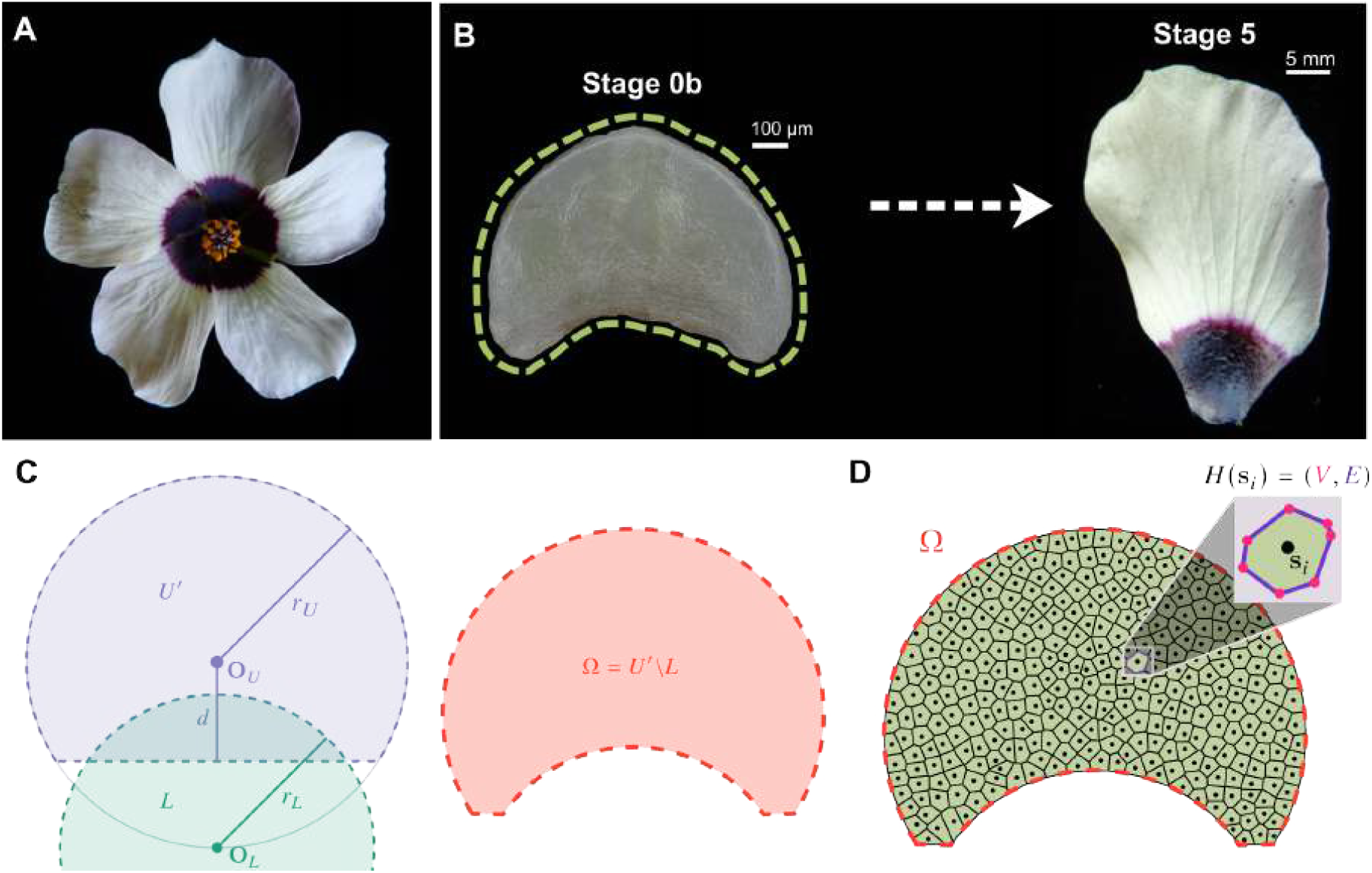
Approximation of the *H. trionum* petal adaxial epidermis tissue shape at developmental Stage 0b. (**A**) The flower of *H. trionum* features a bullseye pattern on its corolla. (**B**) Cell differentiation across the adaxial epidermis of the *H. trionum* petal primordium leads to bullseye pattern formation during petal development. As the petal is likely pre-patterned at an early developmental stage, we created a cell-based developmental model of the Stage 0b petal. (**C**) The Stage 0b tissue shape is approximated by the difference between the major segment of a larger upper circle *U* ^′^ and a smaller lower circle *L*. (**D**) The resulting petal tissue domain Ω is discretised into *N*_*C*_ = 320 cells through Voronoi tessellation. Each Voronoi cell and its Voronoi edges are drawn along with a dot representing its site (see Methods).

**Figure S2.**
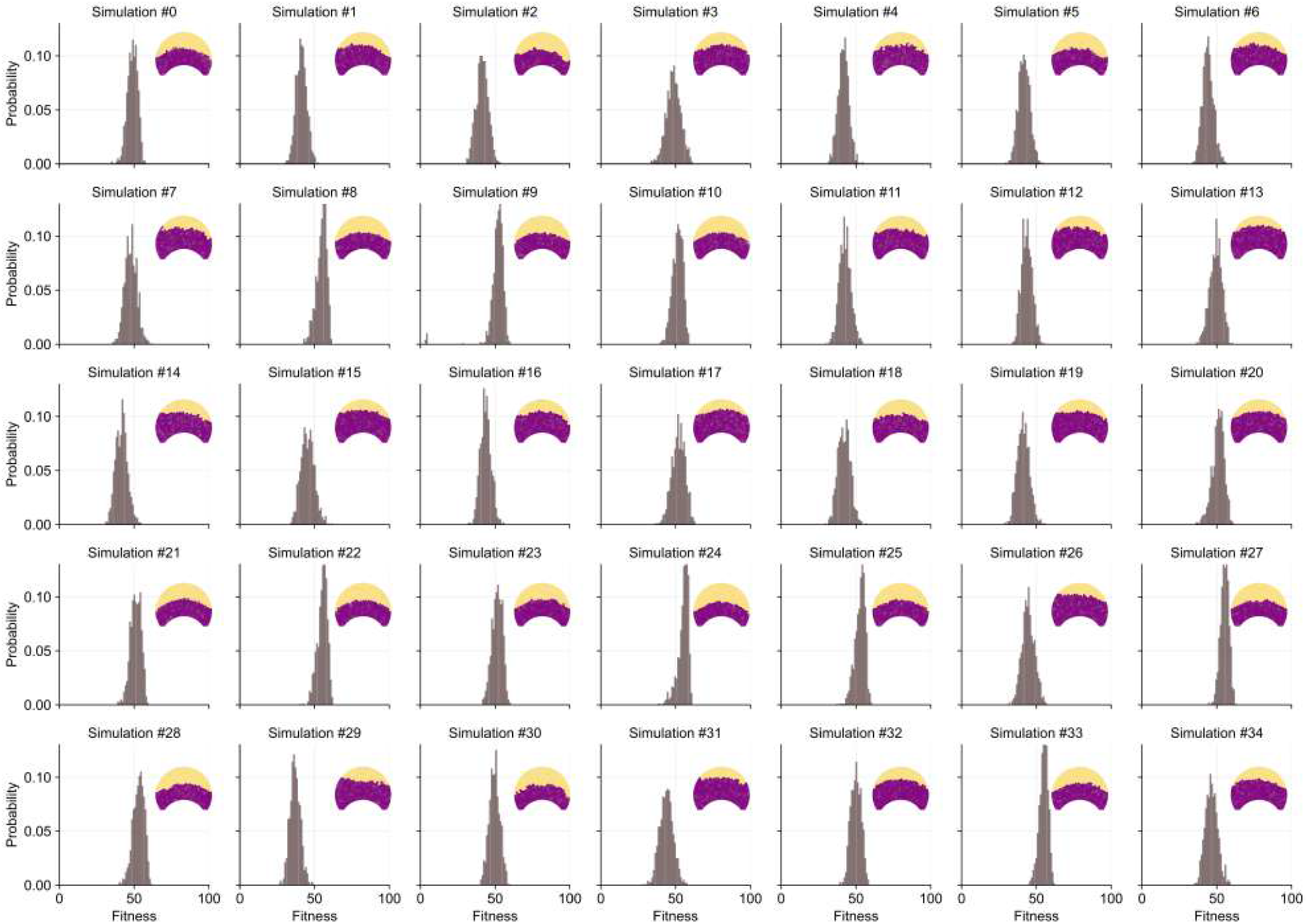
All 35 populations evolved GRNs that robustly generate bullseye patterns. For each simulation, we tested the developmental robustness of the fittest evolved individual by repeating its development 1000 times, each time using a different tissue morphology (see Methods).

**Figure S3.**
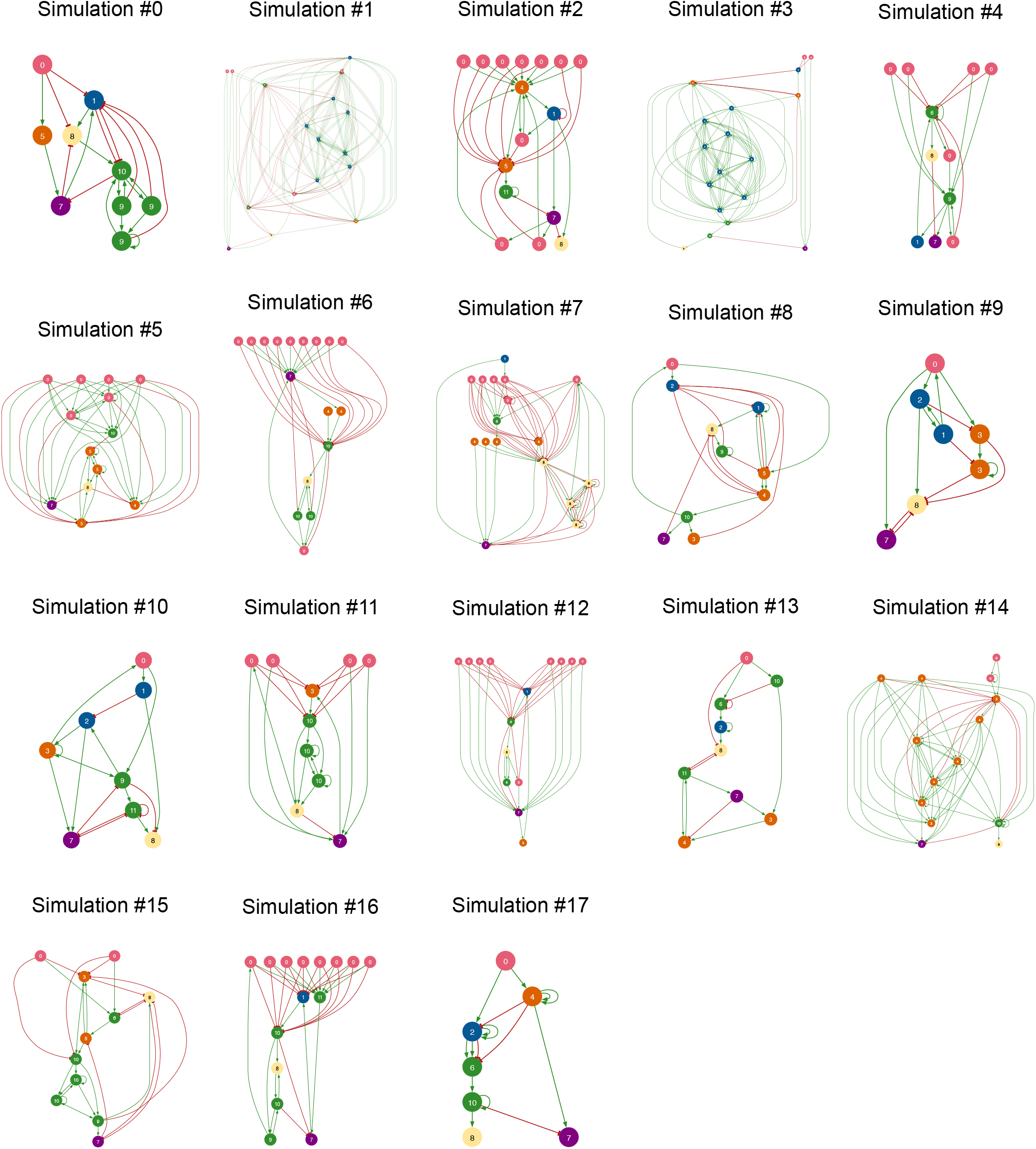
Pruned GRN of the fittest individual in the final generation for simulations 0 to 17.

**Figure S4.**
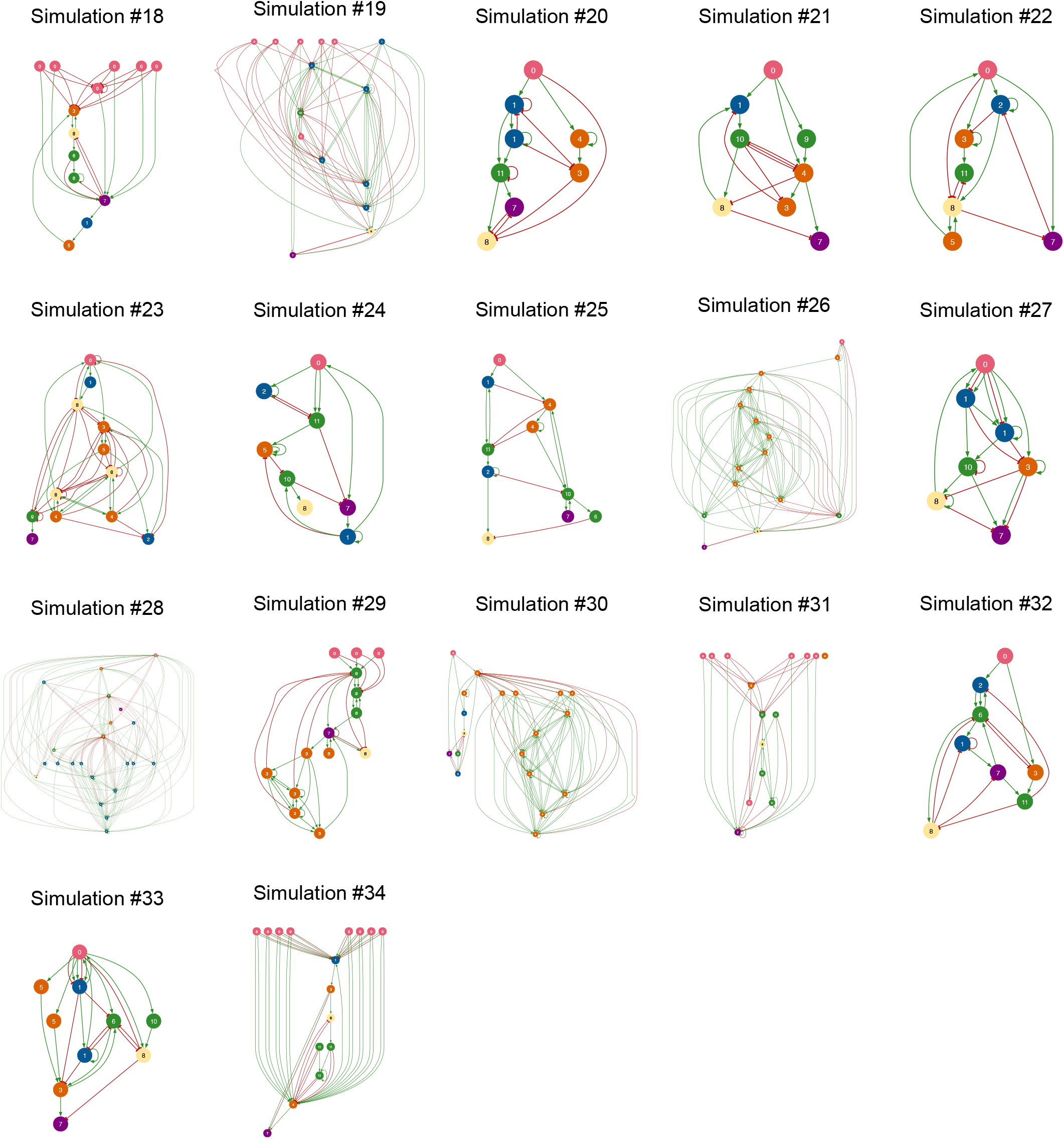
Pruned GRN of the fittest individual in the final generation for simulations 18 to 34.

**Figure S5.**
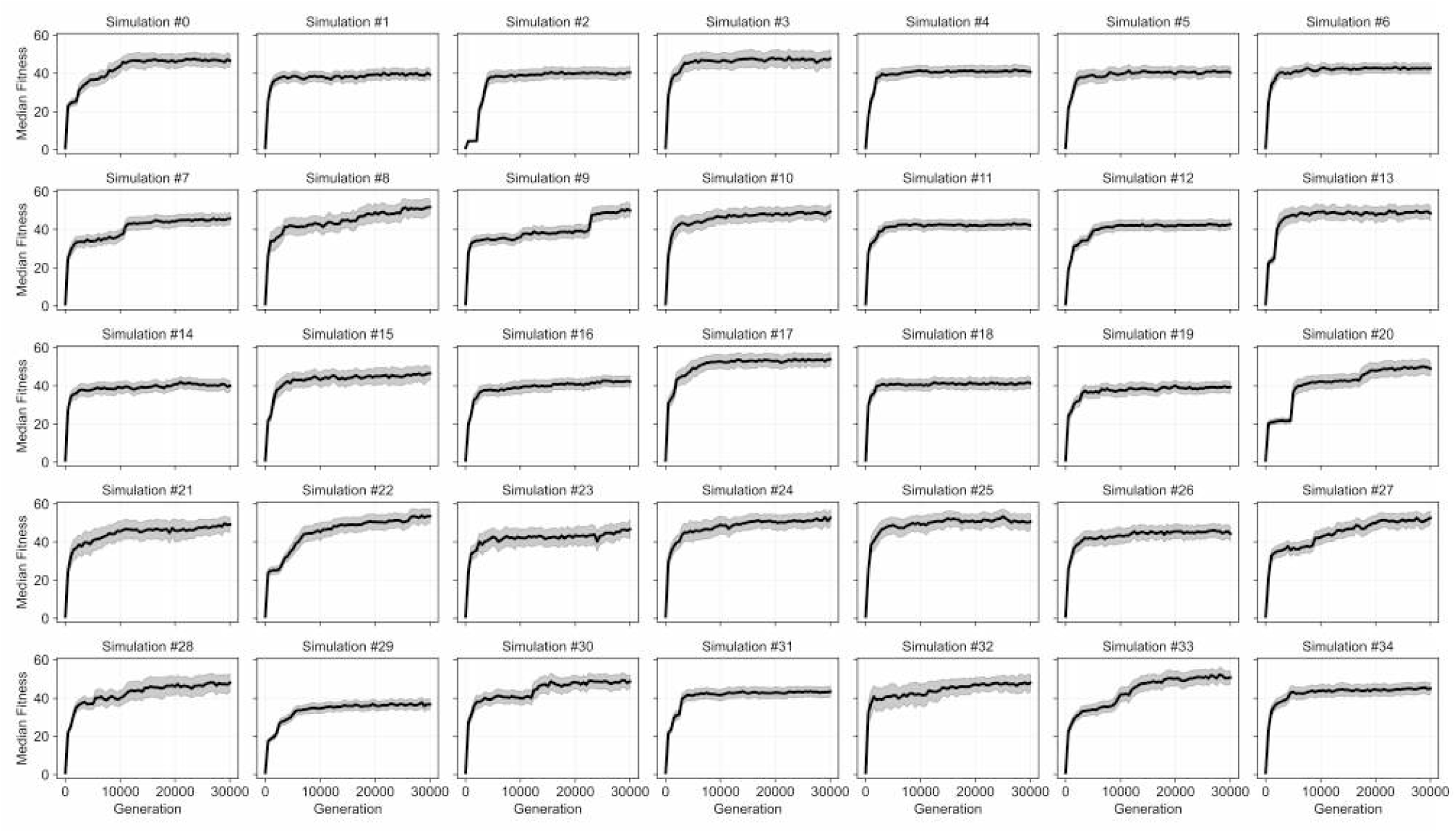
Evolutionary dynamics of population median fitness (solid black line) for all 35 simulations. The median fitness of the population along with the interquartile range (IQR, grey shaded region) is shown for each simulation.

**Figure S6.**
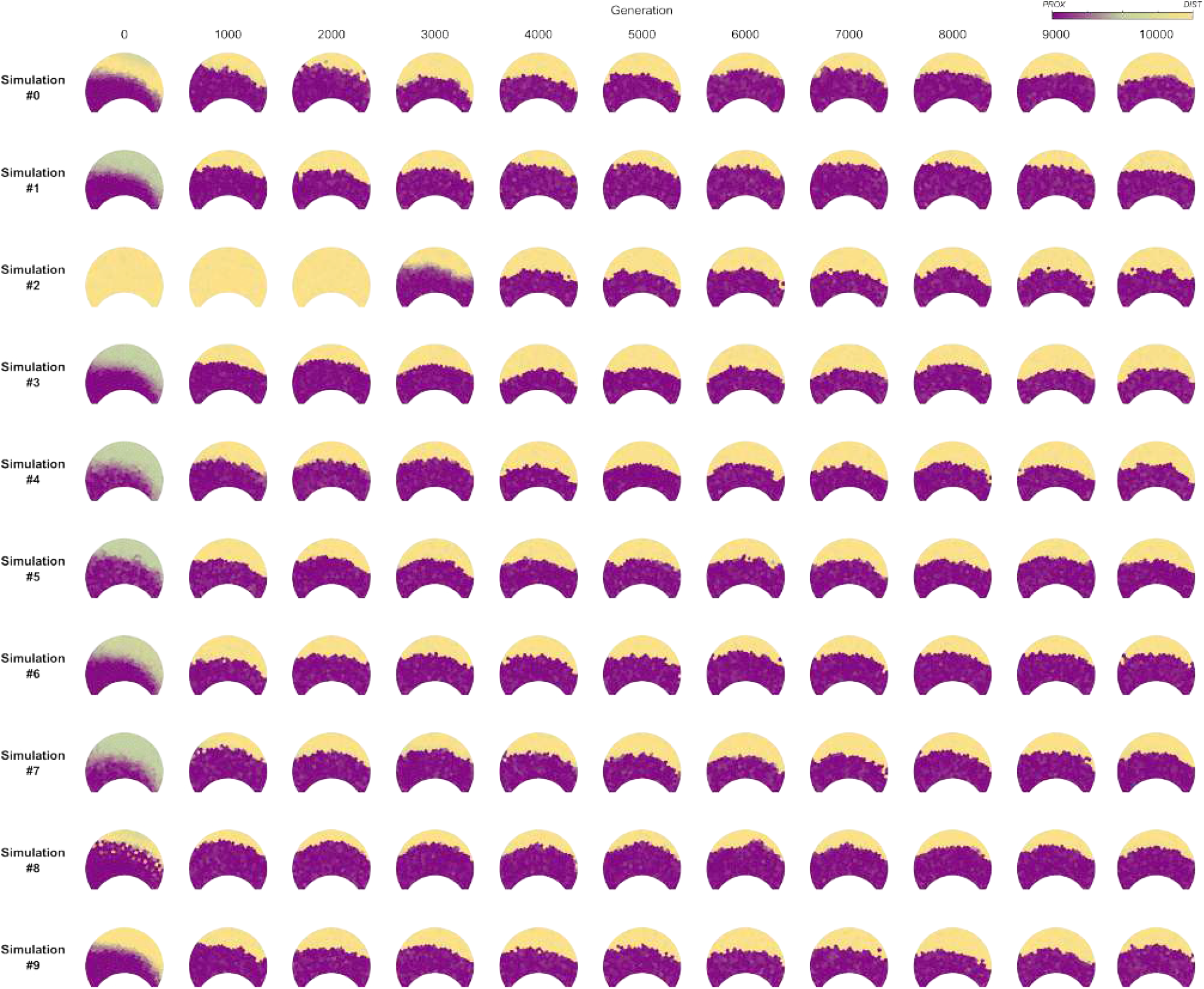
Phenotype evolution of early generations (generations 0 until 10,000) of 10 simulations.

**Figure S7.**
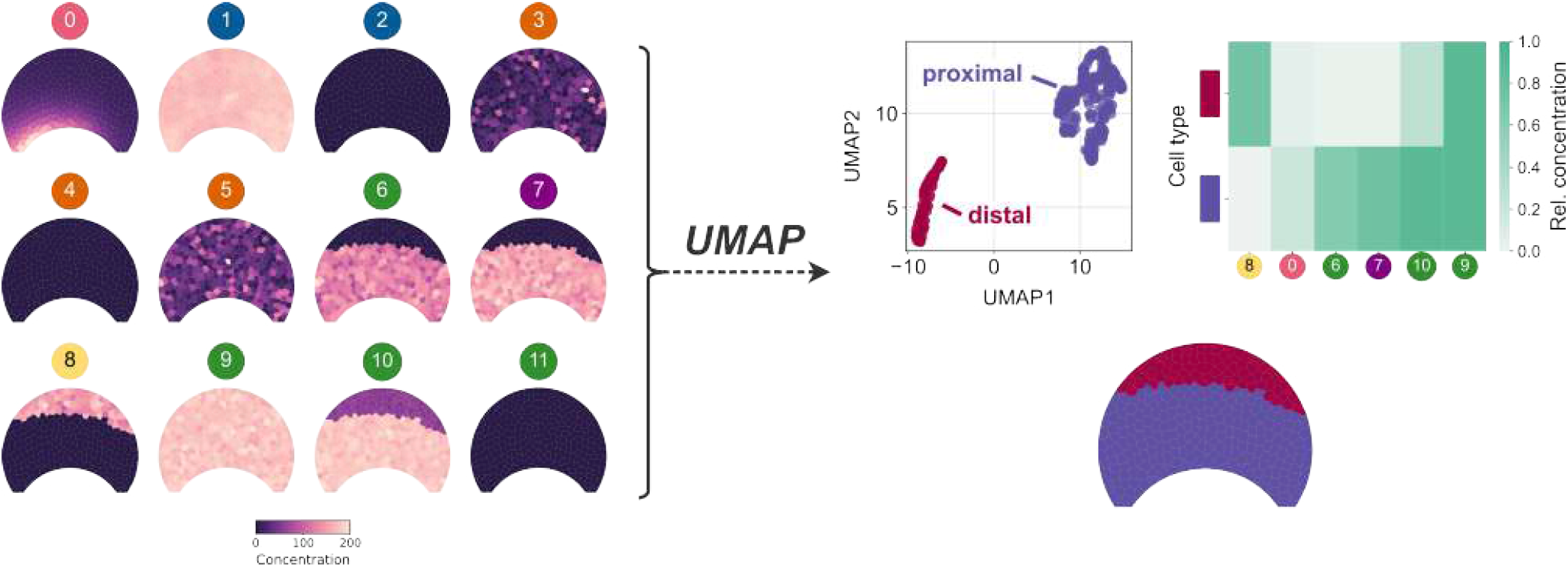
UMAP dimensionality reduction followed by HDBSCAN clustering reveals cell types across petal. UMAP dimensionality reduction was applied to normalized protein concentrations after filtering out low-relevance proteins. HDBSCAN clustering was then performed to identify cell types, which are visualized in a heatmap and across the petal’s epidermal cells. Refer to Methods: Cell Types and Dimensionality Reduction and Clustering for more information on this analysis.

**Figure S8.**
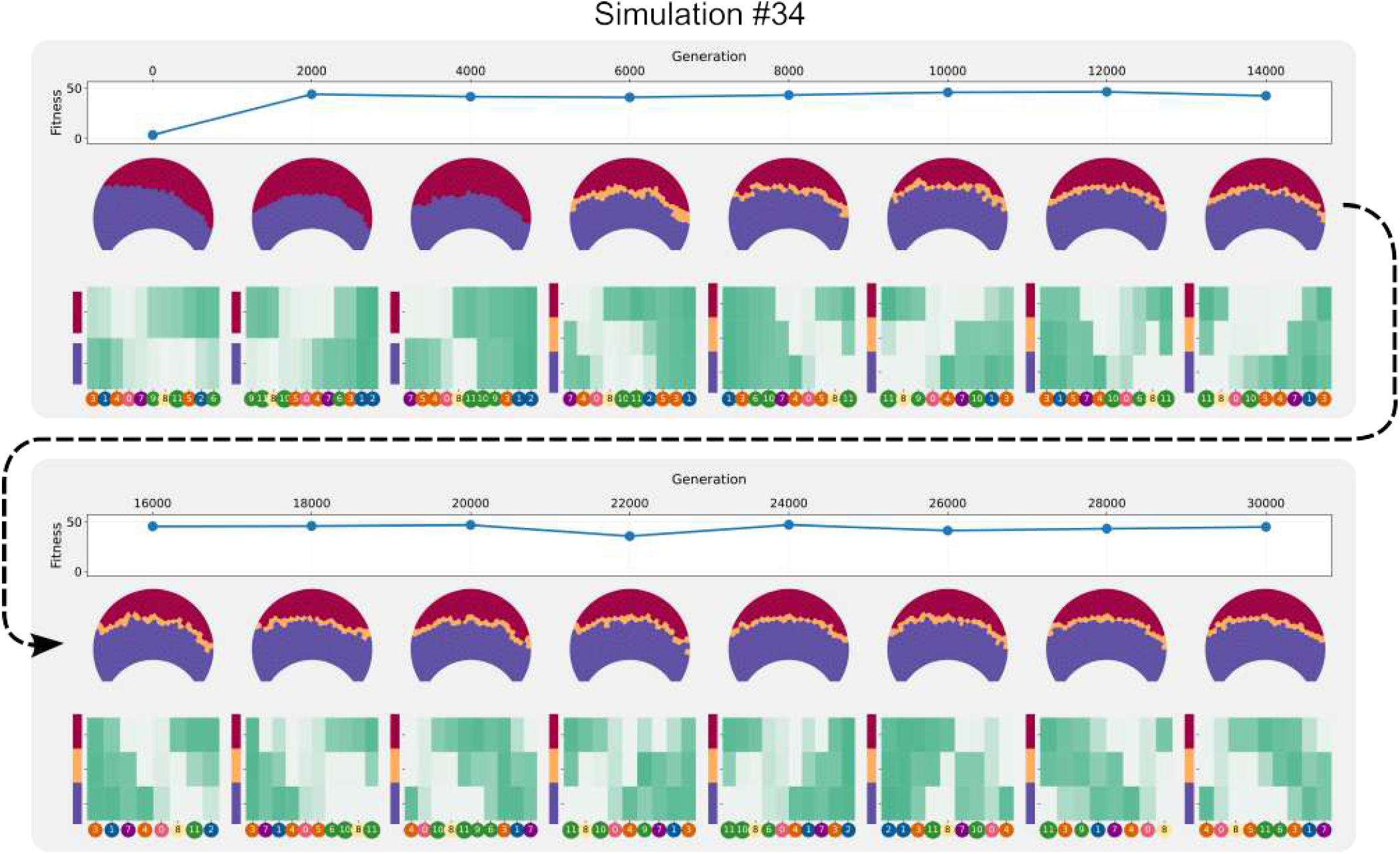
Evolutionary trajectory of the fittest individual in the final population of Simulation #34. Shown are the ancestors of the individual with highest fitness in generation 30,000, together with their fitness and cell types.

**Figure S9.**
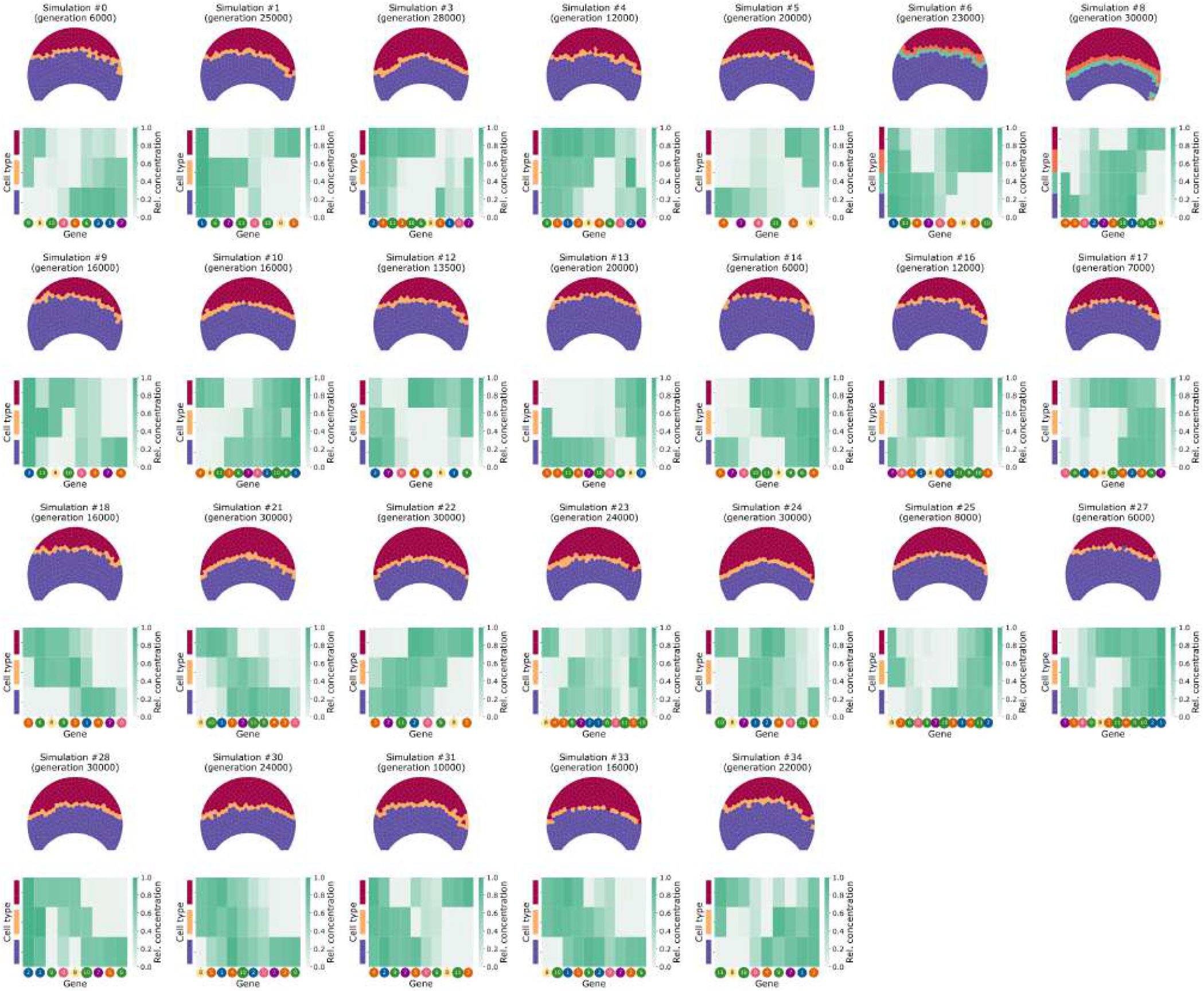
Overview of all noisy simulations that evolved boundary cell types. For each simulation, a single sample from one generation along the ancestral lineage is shown. Bullseye boundary cell types typically appeared across multiple generations in most simulations (Figure 5).

**Figure S10.**
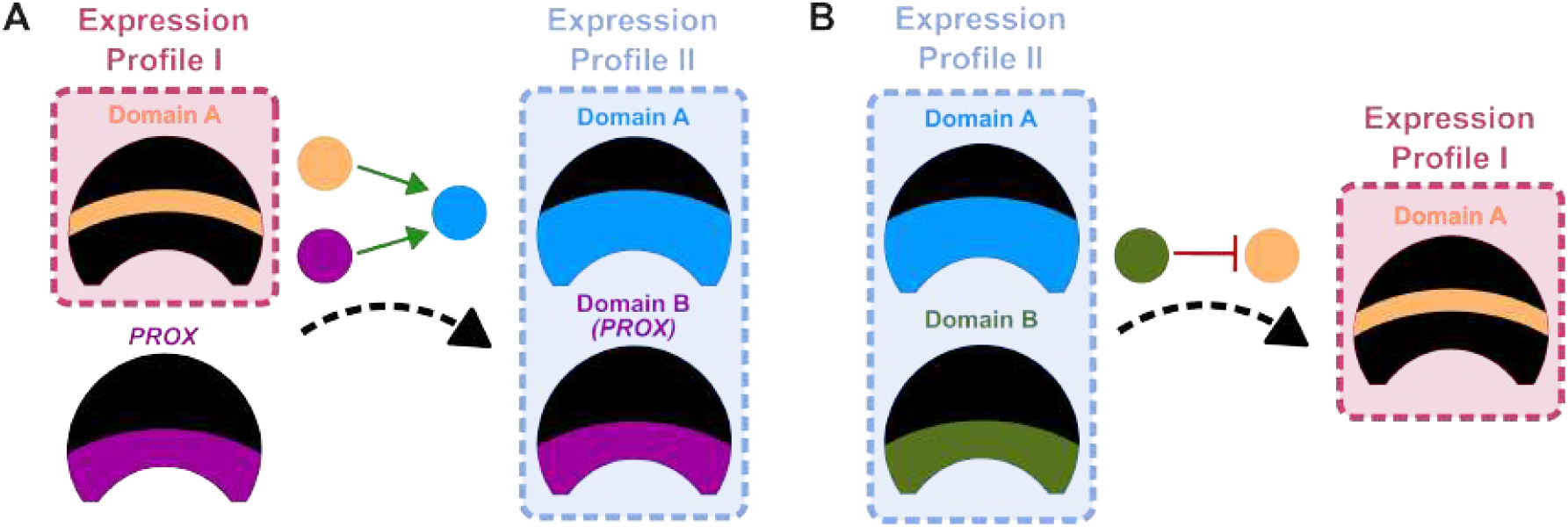
Each boundary expression profile may lead to the emergence of the other. (**A**) Expression profile I can lead to expression profile II by activation of a gene by both the *PROX* and boundary gene. (**B**) Expression profile II can lead to expression profile I when the gene expressed in the smaller bullseye domain (Domain B) inhibits the gene expressed in the larger bullseye domain (Domain A).

**Figure S11.**
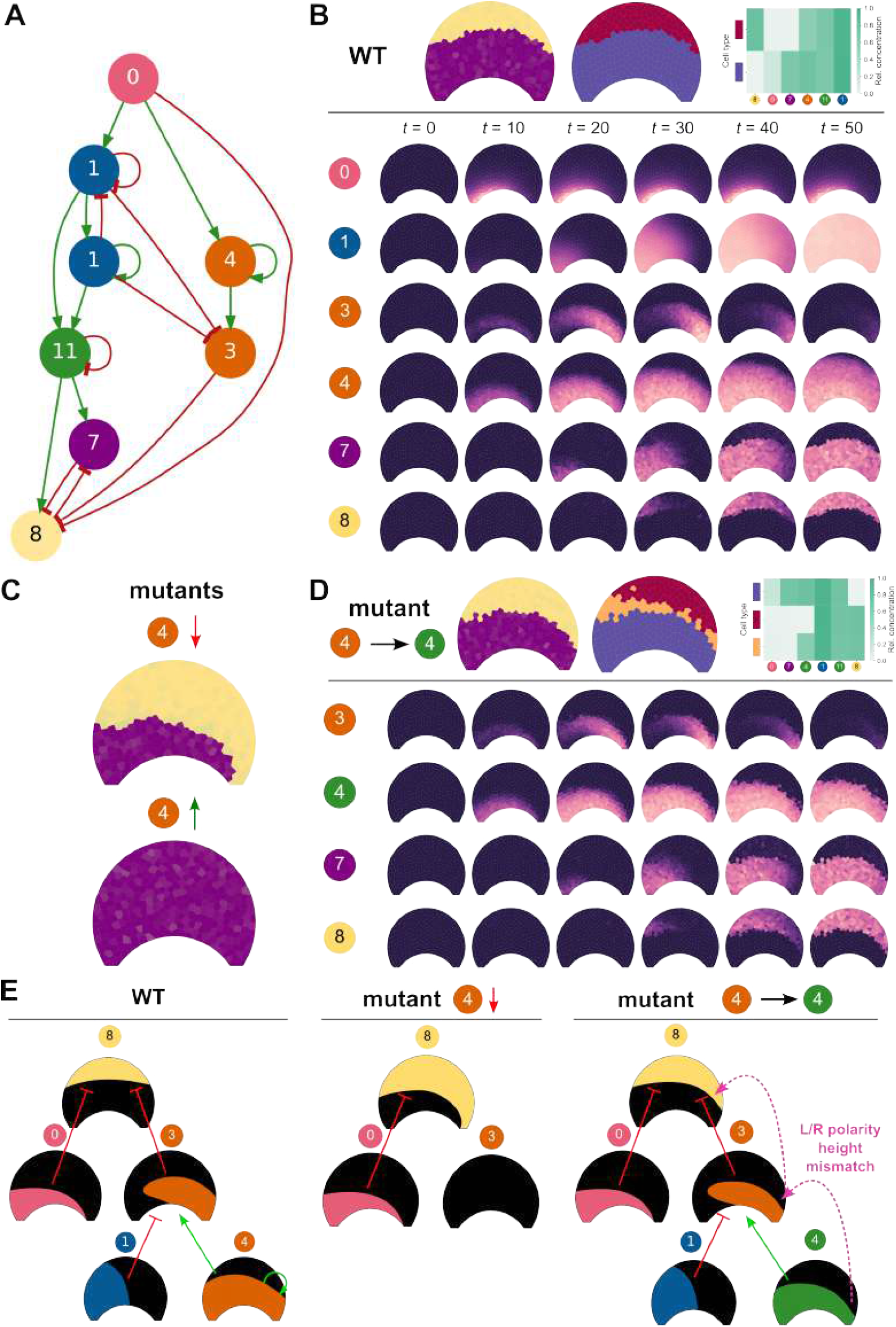
Mechanism of symmetric bullseye patterning without boundary cell type. (**A**) Pruned GRN of an individual from a representative simulation with no boundary cell type (Simulation #20) at generation 30,000. (**B**) Wild-type phenotype and early patterning dynamics. In this GRN, genes 1 and 4 cooperate to establish a temporal right-polarity expression pattern in gene 3, balancing the left-polarity signal in gene 0. Genes 0 and 3 then integrate into gene 8 to produce a symmetric distal bullseye pattern. (**C**) Phenotypic effects of gene 4 knockout and overexpression, resulting in bullseye asymmetry and bullseye loss, respectively. (**D**) Mutant in which gene 4 is converted from a cell-cell communication gene to a TF, with early patterning dynamics shown for downstream genes. (**E**) Cartoon summarising how bilateral symmetry is established in this GRN. In wild type (WT), genes 1 and 4 establish right-polarity expression in gene 3, complementing the left-polarity signal in gene 0 to produce symmetric distal bullseye expression in gene 8. Knockout of gene 4 prevents gene 3 activation, leaving the right side of the bullseye unfilled. When gene 4 is converted to a TF, its self-activation range is reduced: gene 3 is still expressed but the length of its expression domain along the proximo-distal petal axis no longer matches that of gene 0, resulting in a slightly asymmetric bullseye.

**Figure S12.**
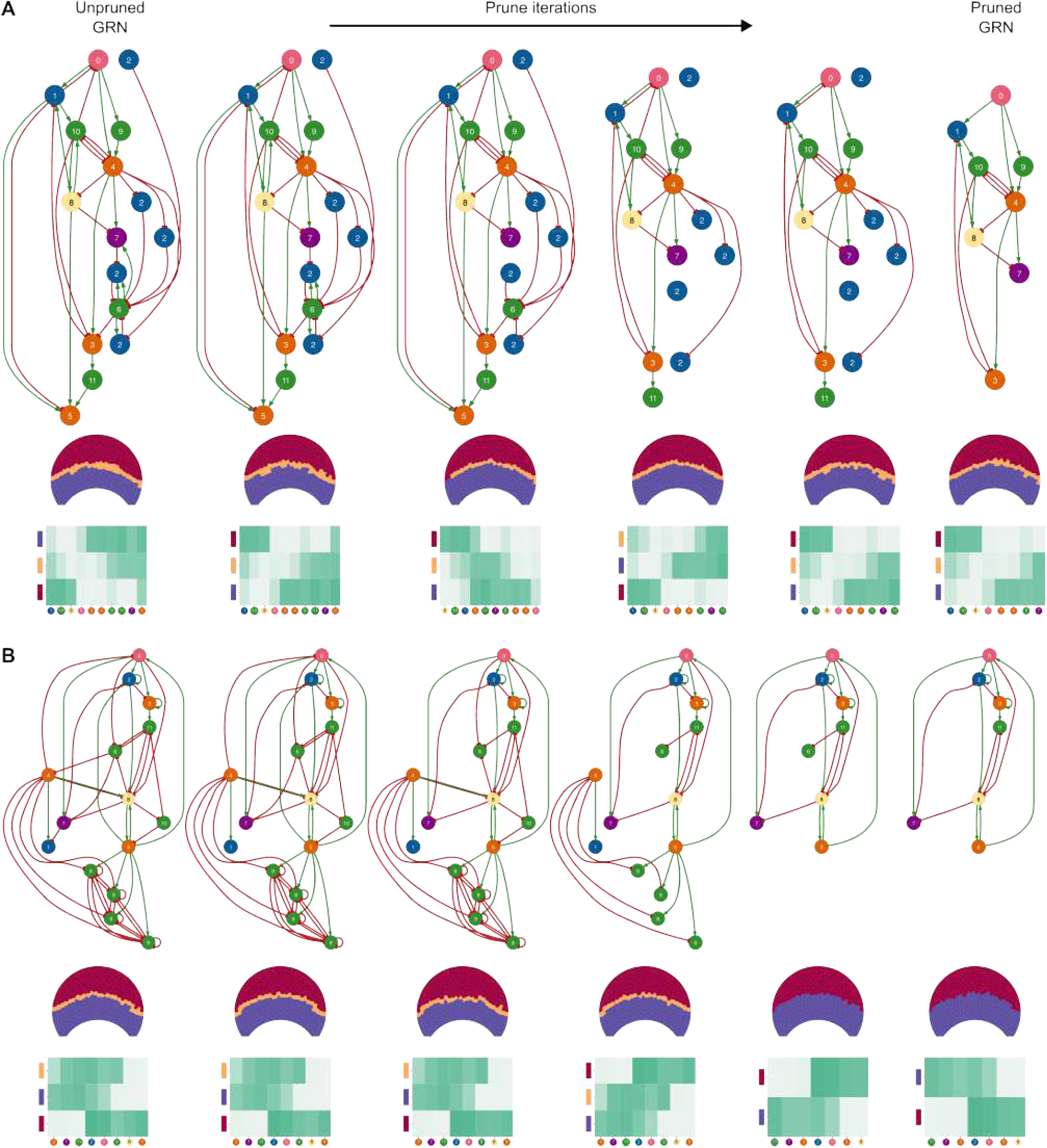
Depiction of the pruning process on two evolved GRNs. When pruning a GRN, we iteratively delete a single gene or interaction and redevelop the pattern 20 times. If the average fitness remains within 3% of the original average fitness score, the deletion is accepted and a next deletion is attempted within the reduced GRN. (**A**) Pruning on a GRN whose boundary cell type persists after pruning. (**B**) Pruning on a GRN whose boundary cell type gets lost after pruning. After pruning, GRNs typically only use a subset of the total 12 gene types available.

**Figure S13.**
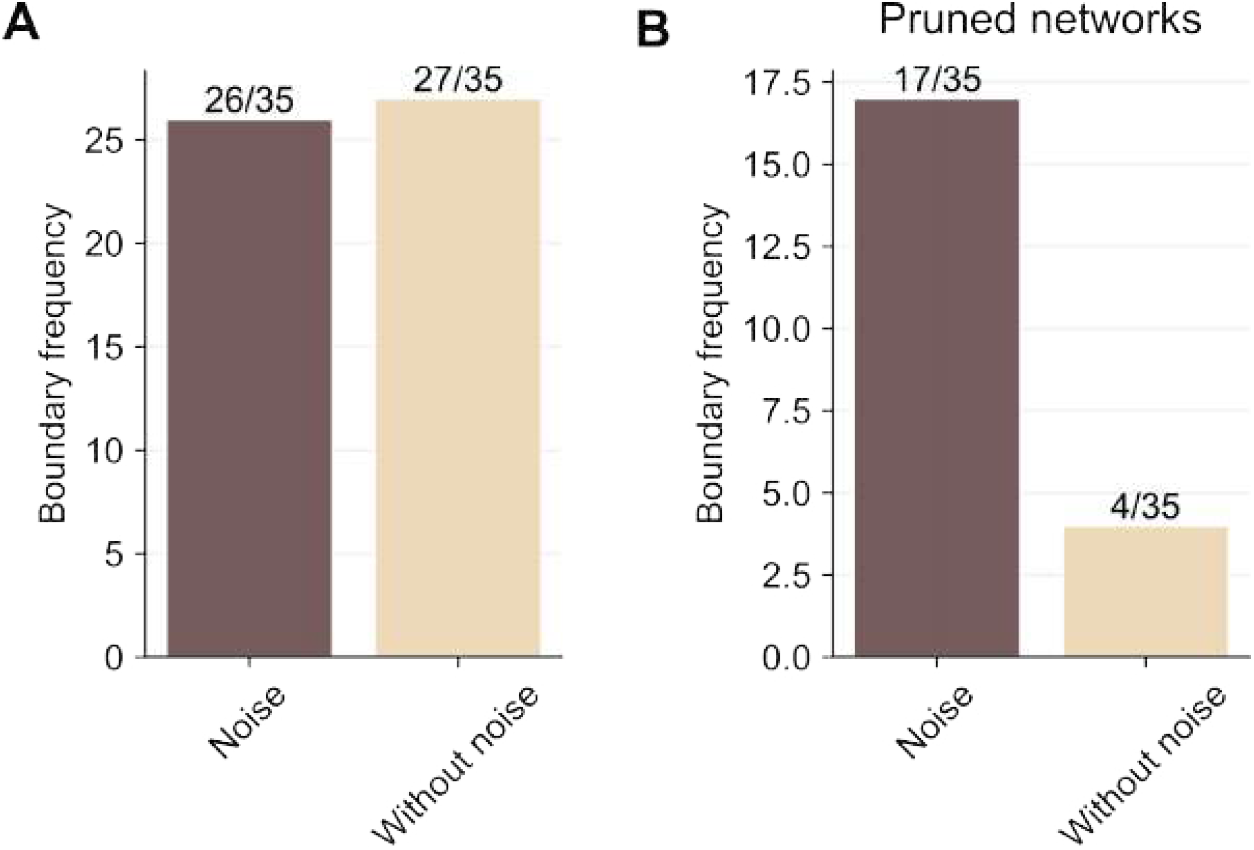
Frequency of evolved boundary cell types across lineages before and after pruning GRNs. (**A**) Number of simulations that evolved a boundary cell type in their lineage, comparing simulations with and without noise. (**B**) Number of networks that have a boundary cell type after pruning (see Methods), comparing simulations with and without noise.

**Figure S14.**
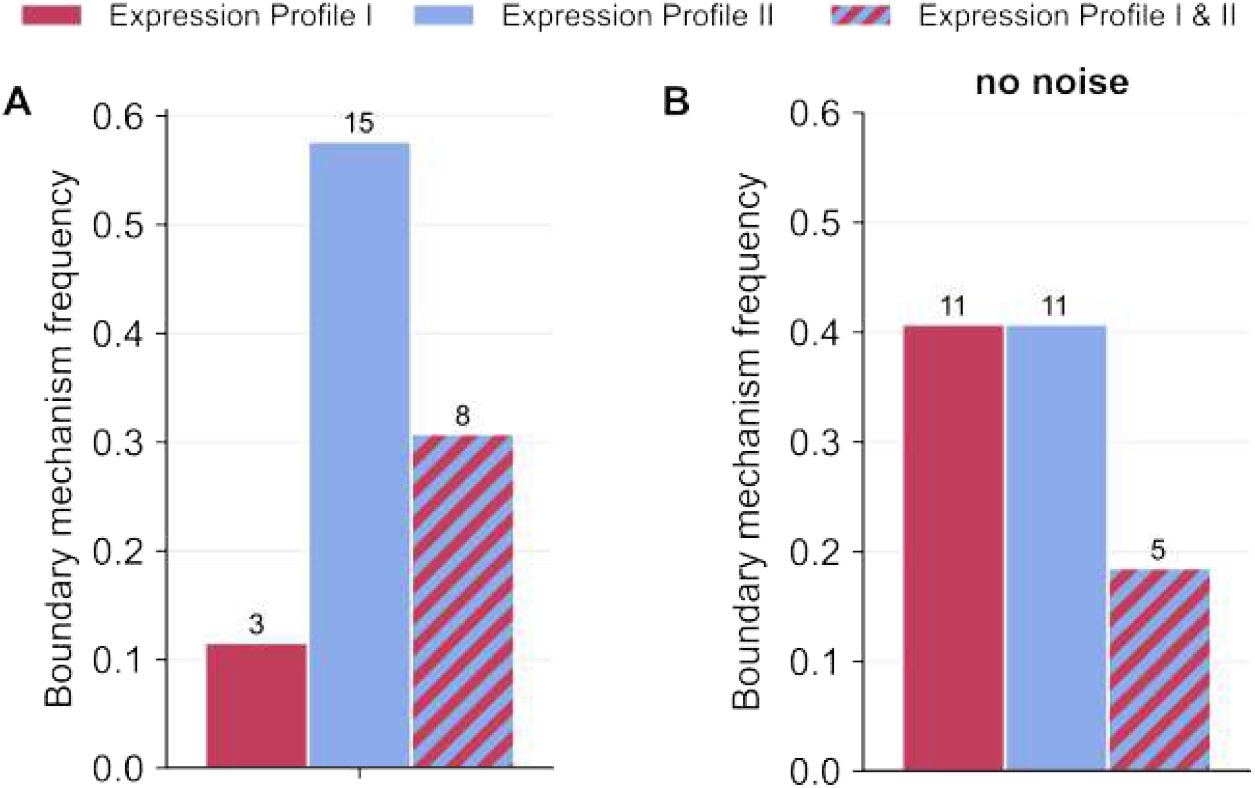
Distribution of boundary expression profiles in the simulations where a boundary cell type appeared in the model with molecular noise (A) and without (B). The absolute number of occurrences for each expression profile is indicated above each bar.

**Figure S15.**
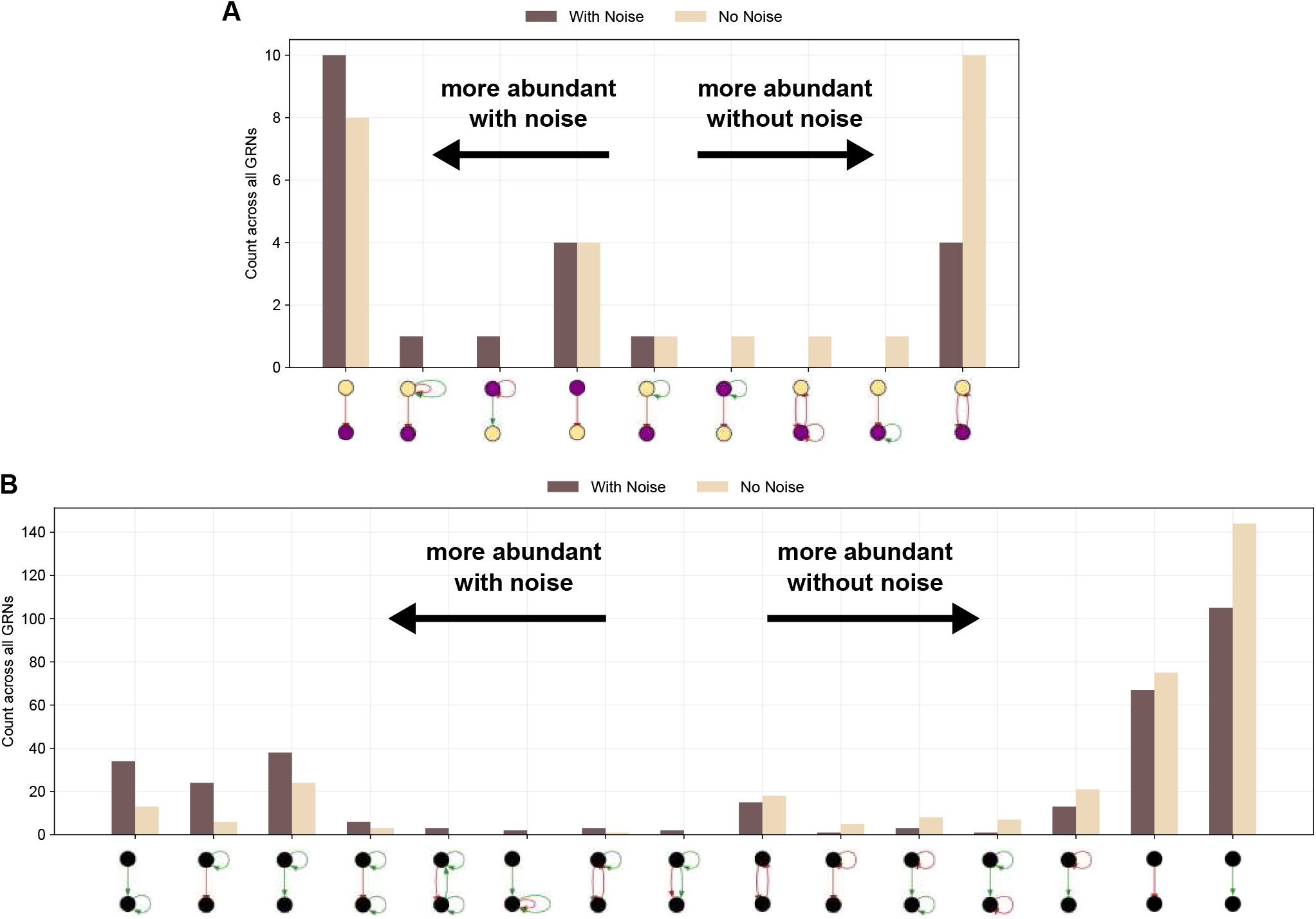
Differences in GRN motifs between simulations with and without noise. (**A**) Differences between 2-motifs containing *DIST* (cream) and *PROX* (purple) genes. (**B**) Differences between 2-motifs of any combination of genes. Note that in simulations with noise, certain motifs are overrepresented due to extensive gene duplications (see Figures S3 and S4).

**Figure S16.**
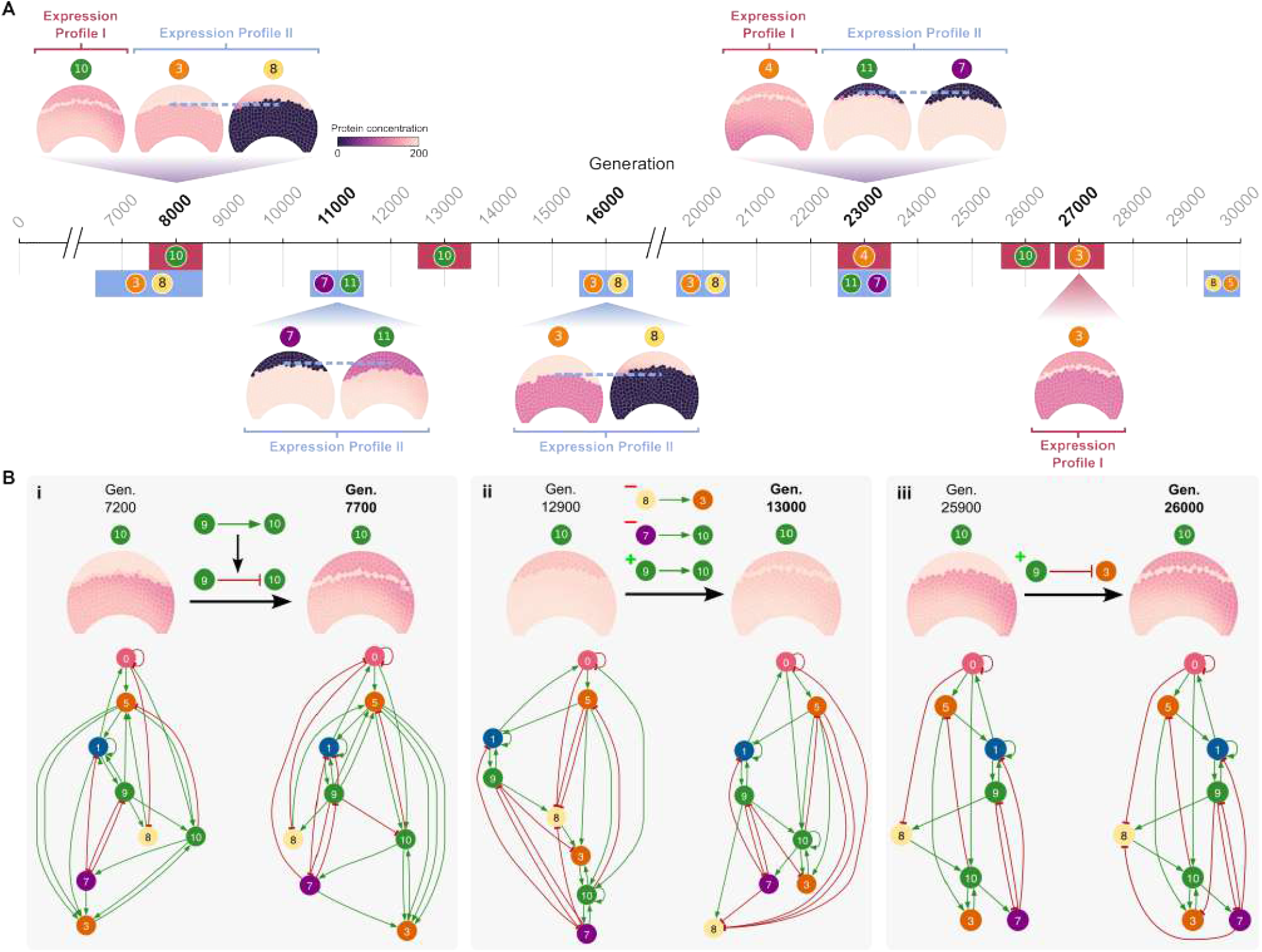
Highly transient evolution of boundary cell type. (**A**) Evolution of boundary cell types in deterministic Simulation #13, illustrating the diversity of boundary mechanisms that evolved within a single simulation. The presence of boundary expression profile I is depicted in pink, while boundary expression profile II is shown in blue. The gene(s) responsible for each emergent boundary cell type are displayed alongside the respective mechanism: for expression profile I, the gene preferentially expressed in the boundary; for expression profile II, the two genes with overlapping expression domains of uneven lengths. Expression patterns of these genes are shown at selected generations. (**B**) For each independent emergence of the boundary cell type mediated by gene 10 (expression profile I), we identify the final mutation(s) responsible. The first appearance at generation 7700 (i) results from a single TFBS weight inversion; the second at generation 13,000 (ii) from three mutations (two TFBS deletions and one TFBS insertion); and the third at generation 27,000 (iii) from a single TFBS insertion. All GRNs shown are unpruned networks with leaf and unreached nodes removed (i.e., genes that do not regulate other genes or are never activated) for visual clarity.

## Supplementary Videos

**Video S1. Representative developmental mechanism which creates a boundary cell type by expression profile I**.

Gene 11 is preferentially expressed in the boundary region, leading to a boundary cell type through expression profile I.

**Video S2. Representative developmental mechanism which creates a boundary cell type by expression profile II**.

Gene 3 and 7 are expressed in proximal domains of different sizes, leading to a boundary cell type through expression profile II.

## Supplementary Tables

**Table S1.**
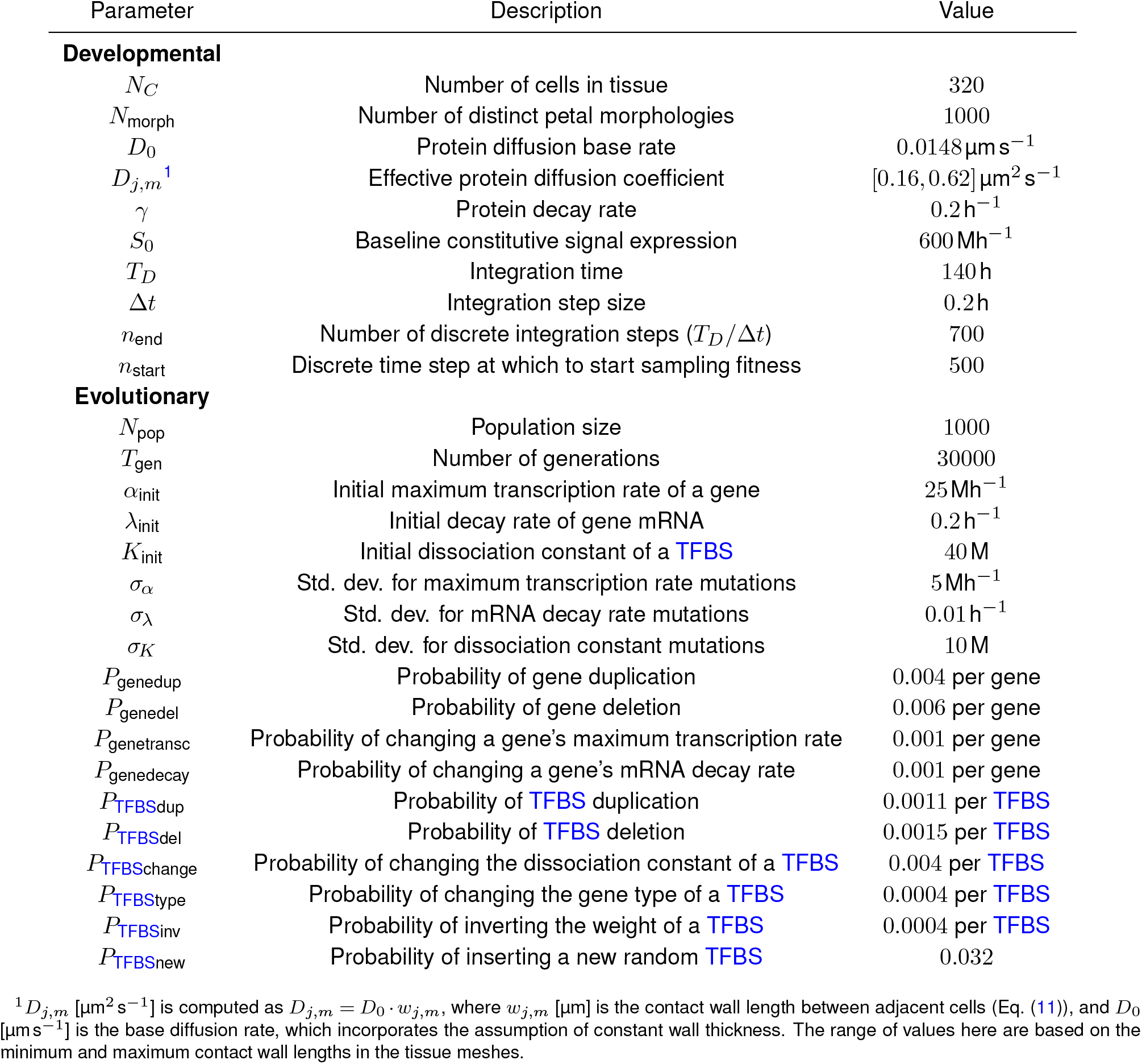
Static parameters for the evolutionary developmental model. Units: M, arbitrary molecular unit; h, developmental time unit (hour); s, second.

**Table S2.**
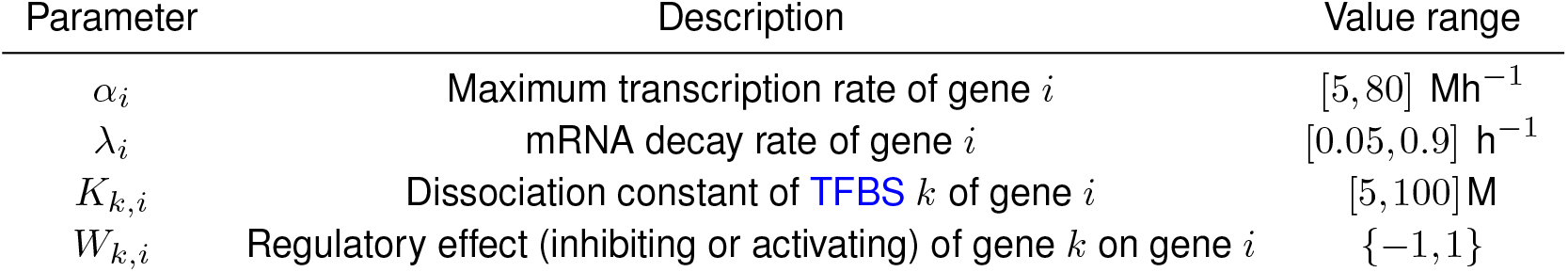
Evolvable parameters and their allowed ranges for the developmental model.

**Table S3. Differential gene expression analysis between proximal and boundary regions of Stage 2 *H. trionum* petal primordia**.

**Table S4. Differential gene expression analysis between distal and boundary regions of Stage 2 *H. trionum* petal primordia**.

**Table S5. Genes preferentially expressed in the petal bullseye boundary region at Stage 2**. Log2 fold Change *<* −1 or *>* 1 and *p*_adj_ *<* 0.05.

## Notes

https://www.ncbi.nlm.nih.gov/datasets/genome/GCA_030270665.1/

https://gitlab.developers.cam.ac.uk/slcu/teamrv/evo-framework/-/tree/paper-2024-stoch-sims

### Competing Interest Statement

The authors have declared no competing interest.

### Summary of Updates

Implement reviewer's comments from Review Commons.

